# Polyglutamine aggregation in the *C. elegans* nervous system is driven by stochastic nucleation events

**DOI:** 10.64898/2026.09.29.755284

**Authors:** Vera van Schijndel, Jorieke Tiggelaar, Myrthe S. E. Franken, Delano Remkes, Maithili Joshi, Deep Sandhu, Klaske Baars, Jesse D. F. de Graaf, Tessa Sinnige

## Abstract

Fibrillar protein aggregation is the hallmark of a wide variety of neurodegenerative diseases. It is still poorly understood how soluble proteins transition to fibrillar aggregates, in particular within the complex environment of cells and tissues. Here, we employ the nematode *Caenorhabditis elegans* to elucidate the molecular mechanisms of polyglutamine aggregation in the nervous system of a multicellular animal. By using fluorescence microscopy in combination with mathematical modelling, we find that polyglutamine aggregation occurs by stochastic nucleation in individual neurons, at a rate that is of the same order of magnitude as previously determined for muscle cells. The J-protein DNAJB6 strongly suppresses polyglutamine aggregation in both tissues, consistent with a mechanism in which aggregation is limited by primary nucleation. We confirm that the same mechanism holds true for a construct including the disease-relevant flanking regions of huntingtin exon 1. Altogether, our data reveal that polyglutamine aggregation occurs spontaneously and largely independently of the cellular context, rather than as a consequence of ageing and the associated decline in protein homeostasis.

## Introduction

Fibrillar protein aggregation is associated with a range of human disorders including Alzheimer’s disease, Parkinson’s disease, type 2 diabetes and Huntington’s disease (Chiti & Dobson, 2017; Louros et al., 2023; Sinnige, 2022). It remains unclear why aggregation is toxic and leads to cell death, and effective treatments are still lacking. Increasing evidence suggests that toxicity does not primarily arise from mature fibrils, but rather from intermediate species formed on- or off-pathway, which can interact with cellular components and disrupt essential cellular processes (Rinauro et al., 2024). To ensure proper protein folding and to counteract proteotoxicity, cells employ a protein homeostasis network including molecular chaperones, protein degradation machinery, and stress response pathways (Hartl et al., 2011; Hipp et al., 2019; Labbadia & Morimoto, 2015). During ageing and under disease conditions, it is thought that the protein homeostasis network becomes overburdened, leading to uncontrolled protein aggregation (Balch et al., 2008; Ben-Zvi et al., 2009; Walther et al., 2015).

Structural studies of fibrils isolated from human brain tissue have provided important insights into the molecular structure of disease-associated aggregates, revealing unique structures in different diseases (Scheres et al., 2023). However, the molecular investigation of protein aggregation in the human brain is restricted to post-mortem analysis, leaving the process of aggregate formation inaccessible to direct observation. *In vitro* aggregation kinetics have revealed the molecular mechanisms underlying the fibrillar protein aggregation of multiple disease-related proteins (Michaels et al., 2018), but lack the complexity of the cellular environment, including active protein homeostasis machineries that are important modifiers of protein aggregation. The aggregation mechanisms of the prion protein and tau have been quantitatively examined in the brains of mice and humans, respectively, by analysing samples taken at different timepoints and fitting the data to mathematical models (Meisl, Hidari, et al., 2021; Meisl, Kurt, et al., 2021). Still, these studies rely on few datapoints and provide limited spatial resolution.

The nematode *Caenorhabditis elegans* provides a uniquely tractable system for studying protein aggregation at (sub-)cellular resolution due to its optical transparency, short lifespan, and stereotypical anatomy. The *C. elegans* nervous system consists of 302 cells, which can be categorised into their functional roles as sensory neurons, interneurons, and motor neurons, and can be further classified based on their neurotransmitter synthesis as e.g. glutamatergic, cholinergic, or dopaminergic (Wang et al., 2024). Transgenic *C. elegans* strains expressing polyglutamine (polyQ) proteins tagged with fluorescent proteins in different tissues have been particularly widely used in the community as a model for studying protein aggregation and its modulation by the protein homeostasis network *in vivo* (see e.g., (Brehme et al., 2014; Brignull et al., 2006; Fawcett et al., 2015; Gutierrez-Garcia et al., 2023; Melentijevic et al., 2017; Morley et al., 2002; Nedosekin et al., 2021; Nollen et al., 2004; Silva et al., 2011; Skibinski & Boyd, 2012)). The aggregation propensity of polyQ proteins can be tuned by their repeat length, where glutamine expansions above ∼35–40 residues drive misfolding and aggregation, in agreement with the pathological threshold in human Huntington’s disease (Chen et al., 2002; DiFiglia et al., 1997; Lieberman et al., 2019).

Previously, we established that the aggregation of polyQ in the body wall muscle cells of *C. elegans* is limited by stochastic nucleation events in individual cells (Sinnige et al., 2021). Once nucleation has occurred, the aggregate grows rapidly to give rise to macroscopically observable inclusions, ultimately leading to each cell acquiring one inclusion. This mechanism is consistent with studies performed in single yeast cells (Kandola et al., 2023) and cultured immobilised rat neurons (Colby et al., 2006), and is also in agreement with the aggregation behaviour of polyQ proteins established in the test tube (Sahoo et al., 2014; Thakur et al., 2009; Torricella et al., 2024; Wagner et al., 2018). However, the intact nervous system presents a far greater level of complexity, comprising different neuronal classes that have been suggested to differ in their vulnerability to protein aggregation (Fu et al., 2018; Kampmann, 2024). Moreover, substantial evidence suggests that misfolded proteins can be transported between cells, where they act as seeds to promote further aggregation, contributing to the spread of protein pathology throughout the brain (Meisl et al., 2020; Peng et al., 2020). Altogether, the challenge remains to quantitatively characterize the progression of protein aggregation across the nervous system at the molecular level, with the aim of identifying the underlying processes that could serve as targets for therapeutic intervention.

Here, we employ *C. elegans* to demonstrate that polyQ aggregation in the nervous system occurs by stochastic nucleation events in the absence of specific neuronal vulnerability. These results imply that polyQ aggregation is spontaneous, rather than a consequence of ageing and the associated decline in protein homeostasis. Our data highlight the chaperone DNAJB6 as a potent inhibitor of polyQ nucleation *in vivo*, although inhibition is incomplete at increasing Q length, and suggest that therapies aimed at lowering the concentration of polyQ fragments will be similarly beneficial.

## Results

### PolyQ aggregation in *C. elegans* neurons is stochastic and occurs at a similar rate as in muscle cells

Previously, we performed kinetic analysis on the aggregation of construct comprising 40 glutamines tagged with yellow fluorescent protein (Q40-YFP) in the *C. elegans* body wall muscle cells, which are large, uniform and easy to visualise by light microscopy (Sinnige et al., 2021). The neurons are much smaller and more heterogeneous in morphology, making it more challenging to identify inclusions in strains expressing polyQ in this tissue. We therefore used confocal microscopy at high magnification to quantify Q40-YFP inclusion numbers in the head region of the animals. We observed that the signal in day 1 adult animals was mostly diffuse, whereas in older animals, multiple bright foci were present (Fig. 1a). These inclusions were absent in Q0 control animals (Fig. S1).

**Figure 1.**
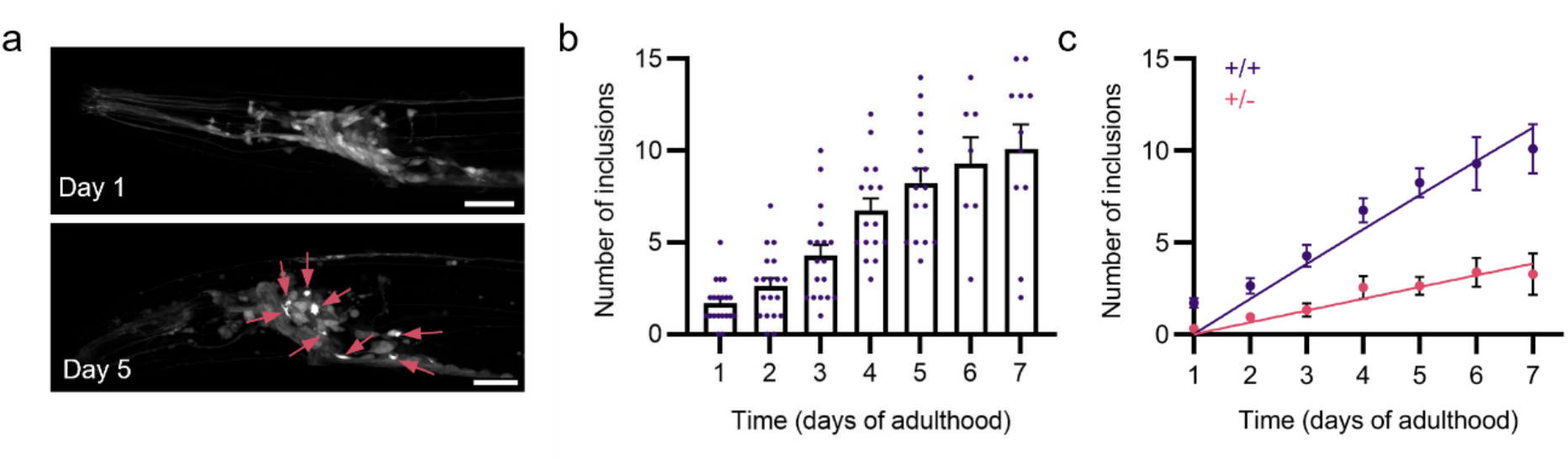
Confocal microscopy reveals that Q40-YFP aggregates through independent nucleation events occurring throughout the nervous system of *C. elegans*. **a** *C. elegans* head region of Q40-YFP animals imaged at day 1 and day 5 of adulthood. Inclusions stand out as bright foci indicated with arrows. Scale bars: 20 µm. **b** Quantification of Q40-YFP inclusion numbers in the head region shows an increase over time. **c** Inclusion numbers for homozygous (+/+) and heterozygous (+/-) animals fitted to a model of independent nucleation events occurring in individual cells. The maximum number of cells was set to 180 and *K*_*cell*_ and *n*_*c*_ were fitted globally. Data represent mean ± SEM (N = 7-20 animals per condition).

We proceeded to monitor the kinetics by which the Q40 inclusions appear over time. Animals from the same batch were monitored from day 1 until day 7 of adulthood, and a steady increase in inclusion numbers was observed (Fig. 1b). The absence of a lag phase is consistent with earlier results in the muscle cells, suggesting that aggregation occurs at a constant rate. However, the plateau of ca. 10 inclusions per animal is much lower than the number of neurons in the head region, which we estimate to be around 180 out of the 302 total neurons in *C. elegans*. This means that only a limited number of neurons in each animal is affected by aggregation during their lifetime. Note that we were unable to collect sufficient data beyond day 7 of adulthood, given the poor health of this strain.

The finding that inclusion numbers are low compared to the number of cells can be explained by two hypotheses: 1) Q40-YFP forms inclusions only in specific neurons, which are more vulnerable to undergoing aggregation than others, or 2) Q40-YFP forms inclusions with similar probability in all neurons, but the nucleation rate is not sufficiently high for all neurons to be affected during the lifespan of the animals. We examined hypothesis 1 by closely inspecting the localisation of the inclusions. Individual *C. elegans* neurons are difficult to discriminate, and their positions are somewhat variable, but the structure of the nervous system is relatively consistent. Inclusions appeared in various regions, including the sensory endings, lateral ganglion, nerve ring, anterior ganglion, ventral and dorsal nerve cords, without displaying a consistent pattern between individual animals (Fig. S2).

To investigate the hypothesis that all neurons can be affected, yet at a low overall aggregation rate, we performed quantitative analysis of the aggregation kinetics. Both homozygous and heterozygous animals were imaged to obtain data at two different protein concentrations, which were fitted to our previously established aggregation model (Fig. 1c). This model assumes that each inclusion arises from a stochastic nucleation event that occurs with equal probability in all cells, ultimately leading to a plateau value that corresponds to the total number of cells (Sinnige et al., 2021). Fixing the plateau value to 180 for the estimated number of neurons imaged in the head region, we obtained a good fit to the experimental data. Global fitting of the reaction order *n*_*c*_ led to a value of 1.58, similarly as reported for *C. elegans* muscle cells (Sinnige et al., 2021).

Since the head region only contains a subset of the animal’s neurons, we asked whether the same stochastic aggregation behaviour extends to the nervous system as a whole. We therefore acquired images along the entire body of the nematode and fitted this dataset using a plateau value of 302, corresponding to the total number of neurons (Fig. 2). Inclusions appeared in all parts of the body, including the nerve cords and the tail, without any obvious regional preference (Fig. 2a, b), consistent with independent nucleation across the nervous system. The stochastic nucleation model gave a similarly good fit to this dataset (Fig. 2c). Altogether, these results suggest that the relatively simple kinetic model developed for body wall muscle cells also holds for Q40-YFP aggregation in the nervous system of *C. elegans*.

**Figure 2.**
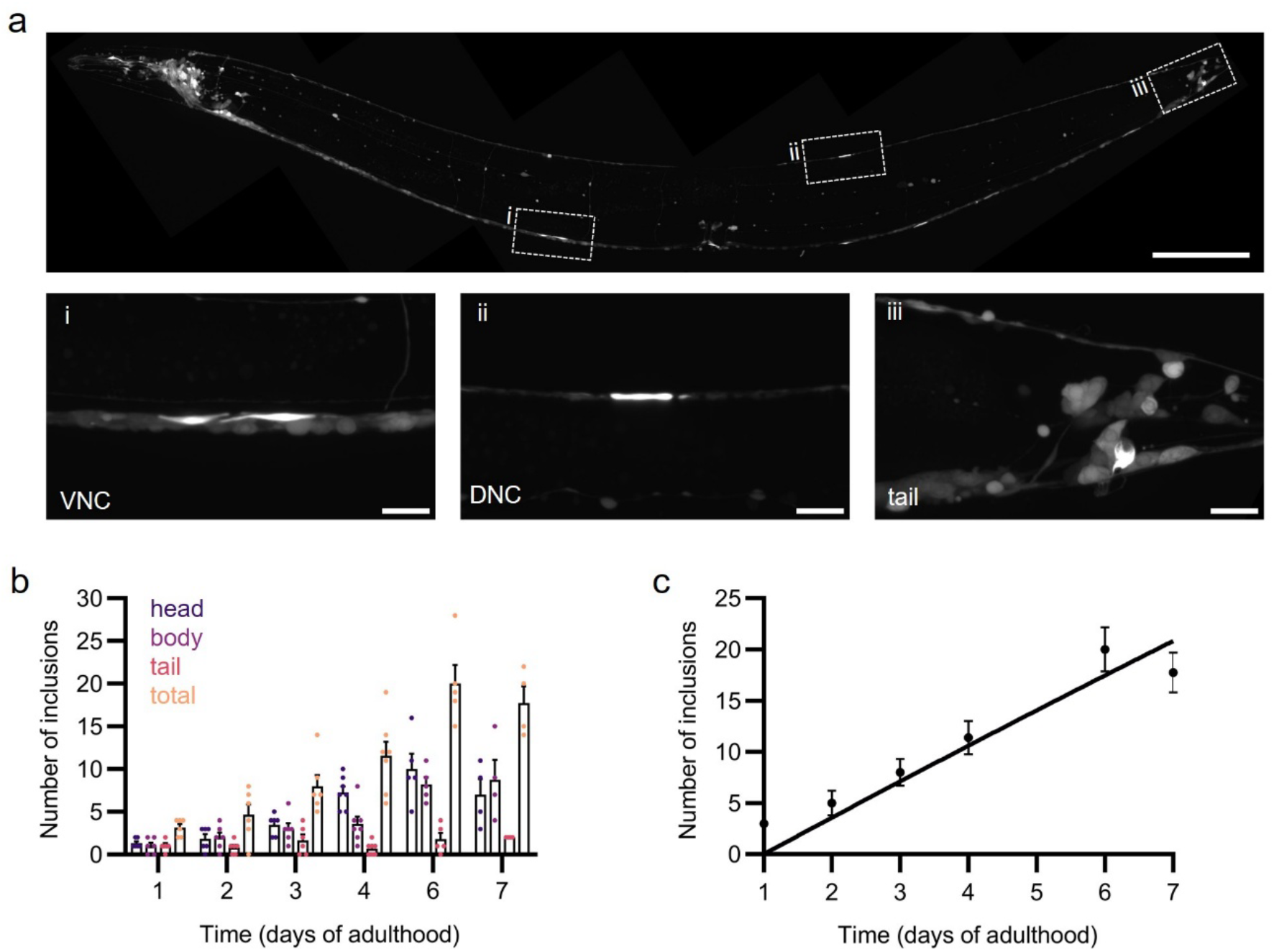
Q40-YFP inclusions appear in all parts of the body. **a** Stitched confocal image of neurons in the whole body of *C. elegans* imaged at day 6 of adulthood. Scale bar: 100 µm. Close-ups display inclusions in i) the ventral nerve cord (VNC), ii) the dorsal nerve cord (DNC), iii) the tail neurons. Scale bars: 10 µm. **b** Quantification of Q40-YFP inclusion numbers in the neurons in the different body parts. **c** Inclusion numbers in the whole body fitted to a model of independent nucleation events occurring in individual cells. The maximum number of cells was set to 302, *n*_*c*_ was set to 1.6 and *K*_*cell*_ was fitted globally. Data represent mean ± SEM (N = 7-10 animals per condition).

In order to extract the nucleation rate constants from the fits to the experimental data, we first quantified the neuronal volume based on confocal images of the YFP control strain. The total fluorescent volume per adult animal was found to be 39 pL, corresponding to an average neuronal cell volume of 131 fL. These values are in reasonable agreement with those determined using the Virtual Worm model (Froehlich et al., 2021) (Table S1). Next, we quantified the amount of Q40-YFP per animal from fluorescence measurements in whole animal lysates (see Materials and Methods). In this way, we were able to determine the average intracellular concentration of Q40-YFP in the *C. elegans* neurons. We applied the same methodology to the *C. elegans* body wall muscle to allow for a quantitative comparison between the two tissues, leading to slightly altered numbers compared to previously published values (Molenkamp et al., 2021; Sinnige et al., 2021) (Table S1). We used the software Amylofit (Meisl et al., 2016) to perform the fitting and obtain the nucleation rate constant, and converted these to the nucleation rate at a fixed protein concentration to allow for a comparison between the different datasets.

At first glance, the nucleation rate in muscle cells appears to greatly exceed that in neurons by a factor of ∼30 (Table 1). However, the actual nucleation rate scales with cell volume at equal protein concentrations, given that more molecules are available to undergo nucleation when their total number increases. According to our calculations, the cell volume of a neuron is ca. 14 times smaller than that of a body wall muscle cell, which may explain the much lower inclusion numbers observed over the same timespan. In fact, converting the nucleation rates to equal volumes, we find that the nucleation rate in neurons is only about 2-fold lower than in the muscle cells (Table 1). This remaining difference may be explained by an excluded volume effect in the body wall muscle cells, where the sarcomere occupies a large fraction of the cytoplasm. This occupancy increases the effective local concentration of polyQ in the available cytoplasmic volume, leading to an increase in the nucleation rate. At the same time, we cannot rule out tissue-specific factors, such as differences in chaperone availability, contributing to the comparatively lower nucleation rate in neurons. Nevertheless, we can conclude that polyQ aggregation in *C. elegans* neurons follows the same mechanism of stochastic nucleation in individual cells with a rate that is of the same order of magnitude as in the muscle cells.

**Table 1.**
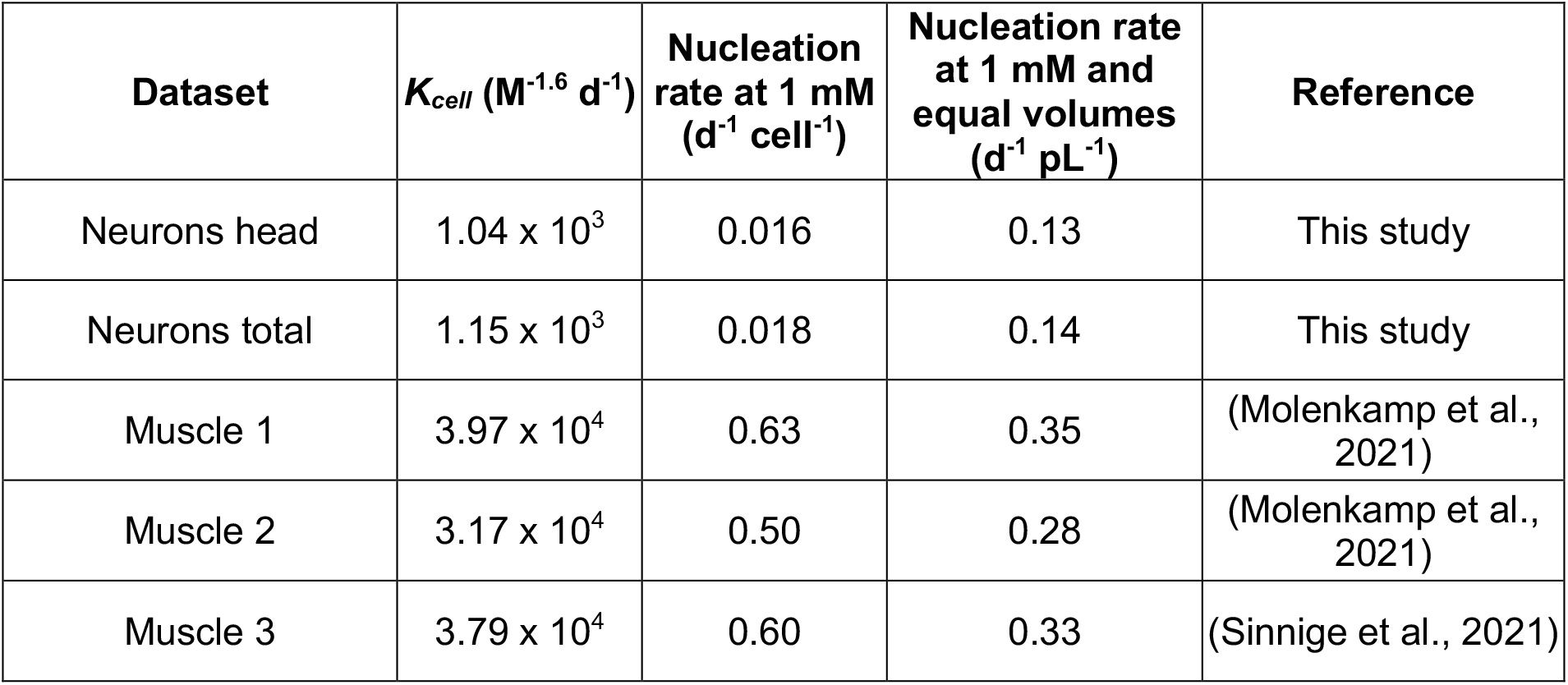
Nucleation rates of Q40-YFP aggregation in *C. elegans* neurons versus body wall muscle cells. The reaction order *n*_*c*_ was fixed to 1.6 for all datasets.

| Dataset | $K_{cell} (M^{-1.6} d^{-1})$ | Nucleation rate at 1 mM ($d^{-1} cell^{-1}$) | Nucleation rate at 1 mM and equal volumes ($d^{-1} pL^{-1}$) | Reference |
| --- | --- | --- | --- | --- |
| Neurons head | $1.04 \times 10^3$ | 0.016 | 0.13 | This study |
| Neurons total | $1.15 \times 10^3$ | 0.018 | 0.14 | This study |
| Muscle 1 | $3.97 \times 10^4$ | 0.63 | 0.35 | (Molenkamp et al., 2021) |
| Muscle 2 | $3.17 \times 10^4$ | 0.50 | 0.28 | (Molenkamp et al., 2021) |
| Muscle 3 | $3.79 \times 10^4$ | 0.60 | 0.33 | (Sinnige et al., 2021) |

### Q67-YFP aggregation affects all *C. elegans* neurons

To further test the prediction that all neurons are ultimately affected by polyQ aggregation, rather than a specific subset, we turned to a longer Q length of 67 residues. In the Q67-YFP expressing strain, virtually all of the protein was found to be already localised to inclusions at day 1 of adulthood (Fig. 3a). Therefore, we set out to image developing animals, starting from a batch of synchronised embryos. The first inclusions appeared very shortly after hatching (t = 12-13 h after a synchronised egg-lay) with the numbers rapidly increasing (Fig. 3b, c). *C. elegans* larvae contain 222 neurons at the time of hatching, and further cell divisions occur during the first three larval stages. The fourth larval stage with a complete nervous system is reached at ∼ 56 h for wild-type animals. Although quantitative analysis of the aggregation kinetics for Q67 is hindered by the increase in both cell numbers and volumes, we observed that the inclusion numbers rose to a plateau value of ca. 300, in agreement with the total number of cells in the completed nervous system (Fig. 3c). Thus, these data support the conclusion that all *C. elegans* neurons ultimately acquire a polyQ inclusion, and that the rate of nucleation limits the number of inclusions that are formed during the lifetime of the animals in the case of Q40.

**Figure 3.**
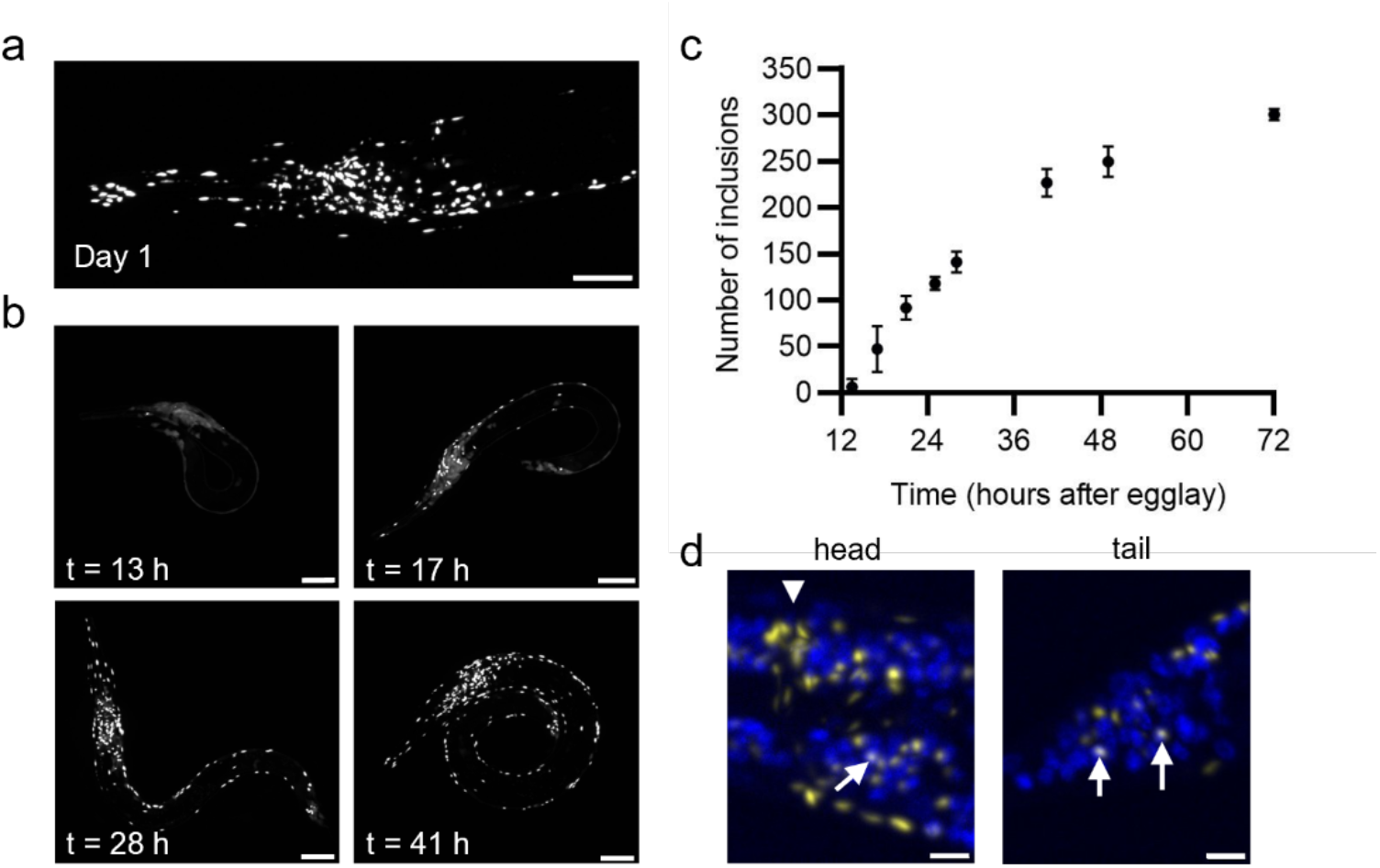
Aggregation of Q67-YFP rapidly takes place in developing *C. elegans* and ultimately affects all neurons. **a** Head region of day 1 adult animal expressing Q67-YFP in the nervous system. Scale bar 20 µm. **b** Confocal images of larvae at different timepoints after a synchronised egg-lay. Scale bars 10 µm. **c** Quantification of average inclusion numbers per animal. Data represent mean ± SEM (N = 7-20 animals per condition). **d** Close-up of a fixed L4-staged animal expressing Q67-YFP (yellow) stained with DAPI (blue). Arrows indicate overlap between Q67-YFP and DAPI (visible as white spots), arrowhead points at the nerve cord. Scale bars: 5 µm.

The larger number of inclusions in the Q67-YFP strain also allowed us to investigate the subcellular localisation of the inclusions (Fig. 3d, Fig. S3). Post-mortem studies of human Huntington’s disease have shown that inclusions may reside in the nucleus and in the cytoplasm of axons and dendrites (DiFiglia et al., 1997; Gutekunst et al., 1999; Herndon et al., 2009). In the *C. elegans* model, we find that the majority of inclusions are cytoplasmic, as we only rarely observe overlap with DAPI staining (arrows in Fig. 3d, Fig. S3b). Inclusions are also found in the nerve ring, which is dense in axons but devoid of cell bodies (Fig. 3d, arrowhead). The distribution of inclusions throughout cytoplasmic regions, and occasionally inside nuclei, suggests that Q67-YFP does not aggregate at a preferential subcellular localisation but that nucleation events occur randomly throughout the cytoplasm and nucleoplasm. Assuming that this result also holds for Q40-YFP, this notion supports our calculations of the nucleation rate based on average total neuronal volumes.

### DNAJB6 effectively inhibits polyQ nucleation in *C. elegans*

The J-protein DNAJB6 has been shown to effectively inhibit polyQ aggregation *in vitro* and *in vivo*. DNAJB6 comprises a serine/threonine-rich region that can form hydrogen bonds with polyQ, thereby suppressing primary nucleation. Furthermore, overexpression of DNAJB6 delays disease onset in a mouse model of Huntington’s disease and reduces the numbers of nuclear aggregates observed at individual timepoints (Kakkar et al., 2016). To shed more light on the molecular mechanism by which DNAJB6 inhibits polyQ aggregation *in vivo*, we expressed mKate2-tagged DNAJB6 in the body wall muscle cells and neurons of the polyQ-expressing *C. elegans* strains and quantitatively monitored the kinetics of Q40-YFP aggregation over time.

Overexpression of DNAJB6 in the nervous system suppressed Q40-YFP aggregation to near completion (Fig. 4a, b). Similar results were obtained in body wall muscle cells, where DNAJB6 overexpression effectively prevented inclusion formation in three strains expressing Q40-YFP at different concentrations (Fig. S4). These results are in line with primary nucleation being the rate-limiting step of polyQ aggregation, and this process being efficiently blocked by the chaperone.

**Figure 4.**
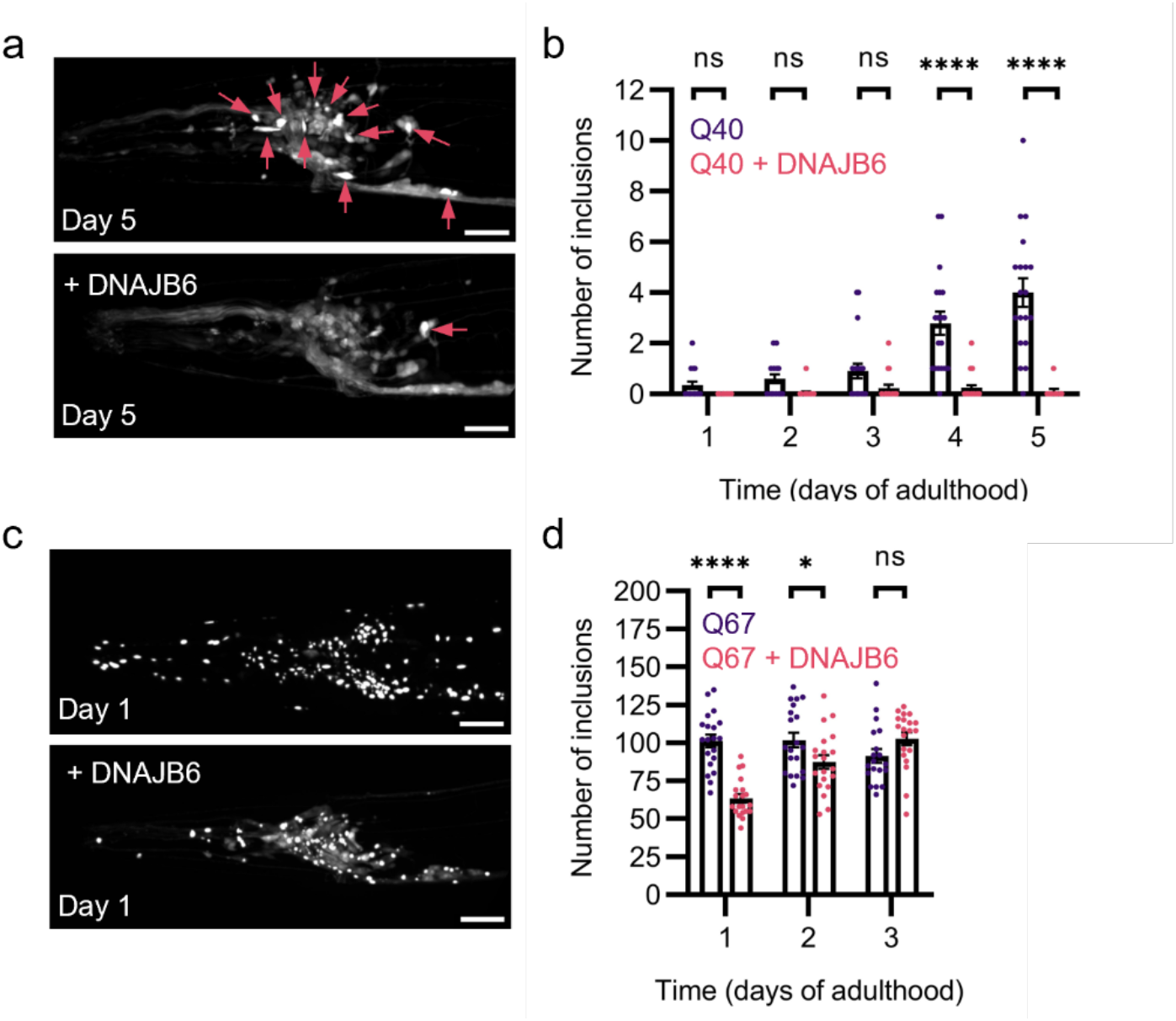
DNAJB6 inhibits primary nucleation in *C. elegans* neurons. **a** Confocal images of the head region of day 5 animals expressing Q40-YFP in neurons under control conditions and upon overexpression of human DNAJB6. Arrows indicate inclusions. Scale bars: 20 µm. **b** Quantification of inclusion numbers in the head region of animals expressing Q40-YFP, compared to animals overexpressing DNAJB6, in which inclusions are largely absent. Data represent mean ± SEM (N = 20 animals per condition). **c** Confocal images of the head region of day 1 animals expressing Q67-YFP in neurons under control conditions and upon overexpression of human DNAJB6. Scale bars: 20 µm. **d** Quantification of inclusion numbers in the head region of animals expressing Q67-YFP, compared to animals overexpressing DNAJB6. Data represent mean ± SEM (N = 20 animals per condition). A two-way ANOVA was performed, followed by Sidak’s multiple comparisons test comparing control to DNAJB6 overexpression on each day. *p* < 0.05 (*), *p* < 0.0001 (****), ns = not significant.

We next asked whether DNAJB6 is also capable of inhibiting inclusions formed by the more aggregation-prone Q67-YFP. We observed that overexpression of DNAJB6 only partially suppressed Q67-YFP inclusion formation as observed in day 1 animals (Fig. 4c, d). Whereas Q67-YFP aggregation is complete at day 1 in control animals (see Fig. 3a), animals overexpressing DNAJB6 display an increase in inclusion numbers up to day 3 (Fig. 4d). These data suggest that in the presence of DNAJB6 also each neuron ultimately acquires an inclusion, but with slower kinetics. Altogether, these results support the notion that DNAJB6 effectively inhibits primary nucleation *in vivo*, similarly as in the test tube (Kakkar et al., 2016; Månsson et al., 2014).

### Expanded huntingtin exon 1 aggregates through a similar mechanism

The polyQ protein used thus far comprises a stretch of glutamines followed by a short linker fused to YFP (Fig. S5), which is a relatively clean system to study the basic principles of aggregation physics. However, this model protein lacks the natural context of disease-related proteins. Therefore, we extended our kinetic framework to huntingtin exon 1 (HTTex1), which is the key pathological protein fragment in Huntington’s disease. HTTex1 contains the N-terminal N17 region that promotes membrane binding and drives oligomerisation, and a C-terminal proline-rich domain that modulates solubility (Fig. S5) (Crick et al., 2013; Hoop et al., 2016).

Confocal imaging of animals expressing HTTex1Q80-YFP in the nervous system revealed bright inclusions, whose number increases with time (Fig. 5a, Fig. S6a). To investigate the aggregation kinetics, we quantified the inclusion numbers in both homozygous and heterozygous HTTex1Q80-YFP animals (Fig. 5b, Fig. S6b). The data fitted well to the independent nucleation model (Fig. 5b, Fig. S6c), suggesting that HTTex1Q80-YFP inclusion formation in neurons is stochastic similar to that of the polyQ model proteins. Global fitting of the reaction order *n*_*c*_ yielded a value of 1.75, which is reasonably close to the value of ∼1.6 obtained for Q40-YFP. Fixing *n*_*c*_ to 1.6 resulted in a similarly good fit to the HTTex1Q80-YFP data, suggesting that the rate-limiting step of nucleation may be the same for the two constructs (Fig. S6c).

**Figure 5.**
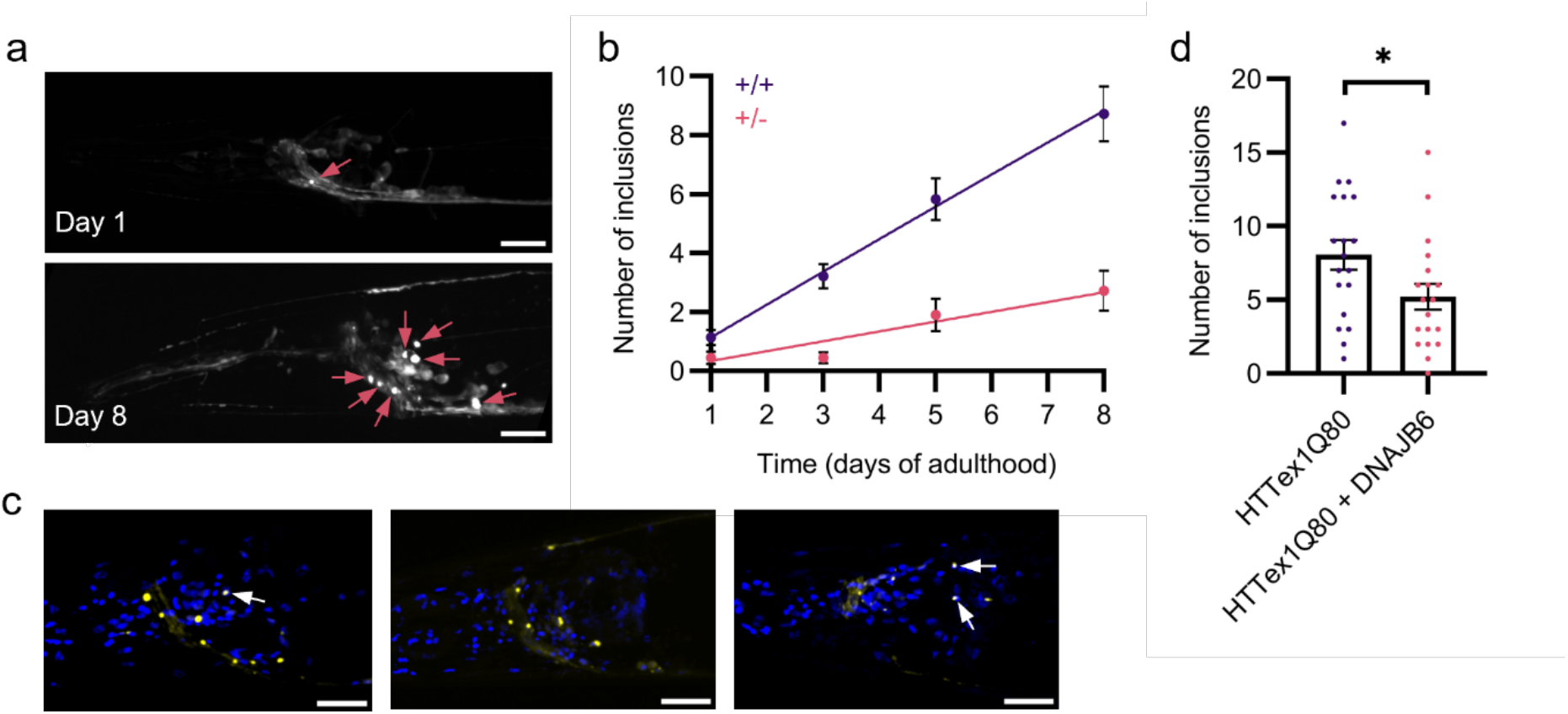
Confocal microscopy reveals that HTTex1Q80-YFP aggregates through independent nucleation events occurring throughout the nervous system of *C. elegans*. **a** *C. elegans* head region of animals expressing HTTex1Q80-YFP imaged at day 1 and day 8 of adulthood. Inclusions stand out as bright foci indicated with arrows. Scale bars: 20 µm. **b** Inclusion numbers for animals homozygous versus heterozygous for HTTex1Q80-YFP, fitted to a model of independent nucleation events occurring in individual cells. The maximum number of cells was set to 180 and *K*_*cell*_ and *n*_*c*_ were fitted globally. Data represent mean ± SEM (N = 18 - 25 animals per condition). **c** Day 8 adult animals expressing HTTex1Q80-YFP (yellow) stained with DAPI (blue). Arrows point at nuclear inclusions. Scale bars: 20 µm. **d** Quantification of inclusion numbers in the head region over time in animals expressing HTTex1Q80-YFP controls compared to animals overexpressing DNAJB6 at day 8 of adulthood. Data represent mean ± SEM (N = 19 animals per condition). *p* < 0.05 (*).

The concentration of HTTex1Q80-YFP was much lower than that of Q40-YFP as determined from the fluorescence images of day 1 animals, differing by a factor of 15 (Fig. S6d). Converting the nucleation rates to equal protein concentrations, we found that the rate of HTTex1Q80-YFP exceeds that of Q40-YFP by a factor of 46 (Table S2), in line with the longer Q length. We verified that the HTTex1Q80-YFP inclusions were mostly cytoplasmic and only occasionally nuclear, similarly as observed for Q67-YFP (Fig. 5c). Furthermore, inclusions were observed in all of the different areas of the nervous system in the head (Fig. S7) as previously identified for Q40-YFP, in agreement with the stochastic nucleation model. Finally, to test whether DNAJB6 is similarly capable of inhibiting nucleation by HTTex1Q80-YFP, we also overexpressed the chaperone in this strain. DNAJB6 overexpression in the nervous system suppressed HTTex1Q80-YFP aggregation, although not to completion (Fig. 5d).

## Discussion

In this study, we used *C. elegans* to investigate the molecular mechanisms underlying polyQ aggregation in the nervous system of a living organism. Whereas aggregation kinetics have been extensively performed in test tube reactions, *in vivo* studies are typically restricted to post-mortem analysis with limited spatial and temporal resolution. By quantifying the numbers of polyQ inclusions over time in living *C. elegans*, we showed that aggregation in neurons is governed by stochastic nucleation events in individual cells.

This molecular mechanism is consistent with earlier findings in single cells and *C. elegans* body wall muscle (Colby et al., 2006; Kandola et al., 2023; Sinnige et al., 2021), demonstrating that the intrinsic biophysical properties of polyQ proteins dominate the aggregation process in diverse cellular backgrounds. Altogether, these studies highlight polyQ aggregation as a spontaneous process resulting from its intrinsic propensity to undergo nucleation, irrespective of the cellular surroundings. Importantly, we find that the rate of nucleation is constant across the adult lifespan of the animal, and does not increase as a function of ageing and the associated decline in protein quality control. These results may be peculiar to polyQ expansions, which are highly insoluble and invariably give rise to disease above the pathological threshold. This contrasts with proteins such as tau or α-synuclein, which form aggregates without being intrinsically very aggregation-prone in sporadic cases of Alzheimer’s and Parkinson’s disease. Aggregation of these proteins is more likely to be actively induced by cellular processes such as post-translational modifications, potentially coupled to aggregate spreading. It will be very interesting to also apply our kinetic approach to these cases.

Cell-non-autonomous effects including spreading are expected to result in a pronounced lag phase in the aggregation curves (Sinnige et al., 2021), which we did not observe for either Q40-YFP or HTTex1Q80-YFP in the nervous system. Cell-to-cell spreading of HTTex1 has been identified in several systems, including *C. elegans* (Kim et al., 2017), *Drosophila* (Babcock & Ganetzky, 2015; Donnelly et al., 2020; Pearce et al., 2015), human cell culture (Herrera et al., 2011), mouse models and human brain (Cicchetti et al., 2014; Pecho-Vrieseling et al., 2014). While our data do not exclude the possibility that aggregated HTTex1 can be transfered between cells, we suggest that the rate of spontaneous nucleation dominates the overall aggregation process when all cells produce approximately equal amounts of the protein. Under human disease conditions this situation likely differs, as detailed below.

We were unable to identify differences in the vulnerability of different neuronal classes to polyQ aggregation. Using a more aggregation-prone strain expressing Q67-YFP, we found that ultimately all neurons acquire an inclusion, as predicted by the stochastic nucleation model. This contrasts with the selective neuronal vulnerability observed in human neurodegenerative diseases, where specific subsets of neurons degenerate while others remain resilient, leading to specific disease patterns (Fu et al., 2018). In Huntington’s disease, medium spiny GABAergic neurons (MSNs) in the striatum are particularly affected, despite mutant HTT being expressed throughout the entire body (Cowan & Raymond, 2006; Morigaki & Goto, 2017). Notably, *C. elegans* possesses a well-defined set of 26 GABAergic neurons (Mclntire et al., 1993), but we did not observe any bias towards this or any other neuronal subtype in the spatial distribution of Q40-YFP or HTTex1Q80-YFP inclusions (Fig. S2, Fig. S7).

Whereas *C. elegans* GABAergic neurons may lack certain features of human MSNs that could explain this result, our data are actually in agreement with the “somatic expansion hypothesis” in Huntington’s disease. This hypothesis states that cells with very long Q lengths due to somatic CAG repeat expansions are the ones that suffer from aggregation and degeneration (Donaldson et al., 2025; Monckton, 2021). Even though the disease occurs above a genetic threshold of 35-40 CAG repeats, the longer, up to hundreds of Q lengths, appear to be associated with pathology. The affected neurons in human Huntington’s disease may thus not be more vulnerable to aggregation by itself, but may have a higher tendency for somatic expansions (Mätlik et al., 2024) and may also produce more of the toxic mutant HTT fragments, for example by incomplete splicing (Neueder et al., 2017). It is conceivable that the aggregated expanded HTT is subsequently transfered to cells carrying the original CAG repeat length, seeding its aggregation which would otherwise happen at much lower probability. Along these lines, exogenous application of HTTex1 fibrils was shown to induce aggregation of a Q15 HTT fragment in *C. elegans* neurons (Skeens et al., 2024), and it would be very interesting to apply our analytical approach to this experimental model as well.

Curiously, we find that the reaction order *n*_*c*_ consistently lies between 1.5 and 2 not only for Q40-YFP in different tissues, but also for HTTex1Q80-YFP. The reaction order informs on the number of molecules involved in the rate-limiting step of the nucleation process, suggesting that more than one molecule is responsible. This result is in agreement with analysis of HTTex1 aggregation in single cells (Colby et al., 2006), but differs from *in vitro* findings. For simple polyQ peptides, monomers have been proposed to undergo nucleation (Chen et al., 2002), whereas HTTex1 was suggested to nucleate from preformed oligomers independently of the monomer concentration (Torricella et al., 2024). Structurally, the nucleus was deduced to comprise a steric zipper formed within a single molecule in the context of yeast cells (Kandola et al., 2023). We cannot exclude that dimerisation of the fluorescent tag contributes to the observed reaction order, although we used a monomeric YFP variant when creating the HTTex1Q80-YFP strain. It may also be possible that a second monomer is required for the nucleus to productively move towards an aggregate in the highly crowded cellular milieu. Further studies using smaller visualisation tags and different sequence permutations may shed more light on this issue.

The stochastic nucleation model implies that once nucleation has occurred, the aggregate will rapidly grow towards a mature inclusion. Therapeutic interventions will thus need to be targeted to this very earliest step of the aggregation process. Our data suggest that lowering the concentration of polyQ protein is an effective means of reducing the aggregation rate, in support of efforts in this direction (Bragg et al., 2026; Bunting et al., 2025; Tabrizi et al., 2019). Alternatively, in agreement with earlier findings *in vitro* and in mouse models (Kakkar et al., 2016), we identify the chaperone DNAJB6 as an inhibitor of polyQ nucleation specifically. However, DNAJB6 failed to completely block the aggregation of Q67-YFP and HTTex1Q80-YFP, whose nucleation probability presumably outcompetes the time that DNAJB6 is bound. Rational designs based on the S/T-rich region of DNAJB6, or completely *de novo* as shown for e.g. amyloid-β and α-synuclein (Sahtoe et al., 2024; Zhang et al., 2026), may lead to more potent inhibitors of polyQ nucleation serving as leads for novel therapeutics.

In conclusion, our work highlights polyQ as a relatively inert protein segment that drives aggregation largely independently of the cellular context. The kinetic framework that we developed is readily extendable to other aggregation-prone proteins relevant for neurodegenerative diseases and may serve as a basis for therapeutic interventions.

## Methods

### *C. elegans* strains and maintenance

*C. elegans* strains used in this study are listed in Table S3. Nematodes were maintained using standard methods on nematode growth media (NGM) seeded with *Escherichia coli* OP50 at 15 or 20 °C (Brenner, 1974). Age-synchronized populations of *C. elegans* were generated by allowing 10 - 15 adult animals to lay eggs on NGM plates for a period of 1-2 h at 20 °C. For long-term experiments, animals were incubated at 20 °C and transferred daily to fresh NGM plates to separate adults from their offspring.

### *C. elegans* strain generation

Human DNAJB6b was codon optimized for *C. elegans* and ordered as a gblock from IDT. The promotor regions of *unc-54, rgef-1* and *rab-3* and the *let-858* 3’UTR were amplified from *C. elegans* genomic DNA (see Table S4 for oligonucleotides used in this study). The *C. elegans* codon optimized mKate2 construct was amplified for N-terminal tagging from the pDD375 plasmid, which was a gift from Bob Goldstein (Addgene plasmid # 91825 ; http://n2t.net/addgene:91825 ; RRID:Addgene_91825). EYFP was amplified from pPD30.38 carrying p*unc-54*::Q40::YFP which was a kind gift from Richard Morimoto. EYFP was converted to mEYFP by introducing the monomerizing A206K mutation via site-directed mutagenesis (Q5 Site-Directed Mutagenesis Kit, NEB). The Huntington exon 1 Q20 and Q80 fragments were amplified from plasmids that were a kind gift from Patrick van der Wel (Poirier et al., 2002). The fragments were recombined in the SalI and SfoI digested pDEST R3-R4 gateway vector using Gibson assembly. Plasmids used in this study are listed in Table S5.

Strains expressing mKate2::DNAJB6b in the neurons or muscle were generated by injecting N2 adult animals with 30 ng/µL pKB06 (muscle) or pJT18 (neurons) together with 70 ng/µL 1 kb generuler (Invitrogen). Extrachromosomal animals were selected based on fluorescence using a Leica M165 FC stereomicroscope with mCherry filter set. Extrachromosomal L4 animals were treated with 30 mJ/cm^2^ UV irradiation using an UVP crosslinker (AnalytikJena). Offspring was singled and screened for integrants using the fluorescent stereomicroscope. Integrated animals were backcrossed 5 times with N2 animals to dilute background mutations caused by UV irradiation, and crossed with Q40-YFP and Q67-YFP expressing animals.

HTTex1Q20-YFP and HTTex1Q80-YFP expressing animals were generated by injecting N2 adults with 30 ng/µL pDS01 and pDS03, respectively, together with 70 ng/µL 1 kb generuler (Invitrogen). HTTex1Q80-YFP animals expressing mKate2::DNAJB6 in the neurons were generated by directly injecting HTTex1Q80-YFP adults with 30 ng/µL pJT18 together with 70 ng/µL 1 kb generuler (Invitrogen).

### Confocal microscopy

For imaging of the *C. elegans* nervous system, animals were immobilised on a 2.5 % agarose pad in a drop of M9 buffer (5.8 g/L Na_2_HPO_4_, 3 g/L KH_2_PO_4_, 0.5 g/L NaCl, 1 g/L NH_4_Cl, 1mM MgSO_4_) supplemented with either 25 mM NaN_3_ or 10 mM levamisole as anaesthetics. Confocal microscopy was performed using a Nikon Eclipse Ti2 body equipped with CSU-W1-T1 spinning disk (Yokogawa, Japan) and a Nikon Plan Apo λD 60x / 1.42, WD 0.15, oil, MRD71670 A Prime BSI sCMOS camera was used to visualize EYFP (excitation 470 nm, emission: 525/50 nm) with 5 % laser power and 100 ms exposure time. Inclusions were counted using the open-source imaging platform ImageJ/FIJI by thresholding the images and using the “analyse particles” plugin on maximum z-projections or the “3D objects counter” plugin on z-stacks in case of overlapping inclusions. Images of the whole worm were stitched using the “Grid/Collection stitching” plugin (Preibisch et al., 2009) before inclusion counting.

Inclusion counting in body wall muscle cells was adapted from the method previously described (Molenkamp et al., 2021). Briefly, 20 animals per well were picked into M9 buffer in a Cellstar flat, clear-bottomed 384-well plate (Greiner, 781091). The animals were immobilised by addition of NaN_3_ at a final concentration of 25 mM 10 minutes before imaging. Confocal microscopy was performed using a Yokogawa CV7000S Cellvoyager equipped with a 2 sCMOS camera, with a 10x objective at excitation and emission wavelengths of 488 nm and 525/50 nm, respectively, using 30 % laser intensity, 25 ms exposure time and 2x2 pixel binning. The wells were imaged in 6 tiles and stitched together with imageJ/FIJI using the “Grid/Collection stitching” plugin (Preibisch et al., 2009). Inclusions were counted using imageJ/FIJI software by thresholding and the “Analyse Particles” plugin.

### DAPI staining

Animals were washed from NGM plates with M9 buffer, put on ice and fixed in 4% paraformaldehyde in phosphate-buffered saline (PBS) pH 7.4 for 20 minutes at room temperature. Pellets were washed three times with PBS + 0.05% Tween-20 and carefully pipetted onto a microscope slide. After briefly drying the suspension, a drop of 10-20 µL of Fluoromount-G with DAPI (Invitrogen) was added and a cover slip was laid on top. Slides were put in the dark at room temperature to dry overnight and stored at -20 °C until imaging.

### EYFP production and purification

EYFP was codon optimized for *E. coli* expression and obtained from IDT as a synthetic DNA fragment. The fragment was assembled into the pET-28b+ vector digested with NotI and SacI using Gibson assembly. The plasmid was transformed and isolated from *E. coli* TOP10 cells and verified using restriction digestion and sequencing.

An overnight culture of BL21 star (DE3) transformed with pJT07 was diluted to an OD600 of 0.1 in 500 mL LB-medium with 50 µg/mL kanamycin and grown for 4 h at 37 °C with shaking at 220 rpm. EYFP expression was induced for 4 h with 0.1 mM IPTG. Bacteria were lysed in a buffer containing 150 mM NaCl, 10 mM imidazole, 3 mM β-mercaptoethanol, 50 mM potassium phosphate buffer pH 7.4 and 1x cOmplete™, EDTA-free Protease Inhibitor Cocktail (Roche) by sonicating for 5 rounds of 30 s with 2 min intervals using the Sonifier 250 cell disruptor with a microtip at 30 %. Cell debris was separated by centrifuging for 30 minutes at 12,000 rpm cooled to 4 °C, filtered through a 0.2 µm filter and loaded upon a 5 mL HisTrap HP His tag protein purification column (Cytvia). Yellow fractions were concentrated to 500 µL using a 10 kDa Amicon centrifugal filter (Millipore) and loaded upon a Superdex 200 increase 10/300 GL gel filtration column (Cytvia) and eluted with 50 mM Tris-HCl pH 7.4 and 150 mM NaCl buffer. Fractions were analysed by SDS PAGE and peak fractions were pooled. The protein concentration was determined using the Nanodrop by measuring absorbance at 280 nm. Purified stocks were aliquoted and stored at -80 °C until use.

### Determining protein concentrations

Protein concentrations were determined by measuring Q40-EYFP fluorescence in cell lysates and comparing it to a calibration curve prepared with purified EYFP. The lysates were prepared by harvesting a large population of day 1 age-synchronized adult animals from NGM plates with M9 buffer. The amount of animals harvested was estimated by counting the number of animals in three 10 µL drops taken from the sample. Pellets were snap frozen in liquid nitrogen and stored at -80 °C until further use.

Pellets were thawed on ice and lysis buffer was added to a final concentration of 50 mM Tris-HCl pH 7.4, 5 mM MgCl_2_, 0.5% Triton-X100, 0.2 mM phenylmethylsulfonyl fluoride (PMSF), 1 µg/mL leupeptin and 1x complete™, Mini Protease Inhibitor Cocktail (Roche). Samples were lyzed 5 for minutes at 30 Hz with a 7 mm stainless steel bead (Qiagen) using the Tissuelyzer II (Qiagen). Debris was removed from the sample by centrifugation for 5 minutes at 3,000 rpm cooled to 4 °C. The amount of total protein per sample was determined using the Pierce BCA protein assay kit (Thermo Fisher). Protein lysates were diluted to equal protein concentrations and the fluorescent intensity of EYFP in the lysates was determined using the Clariostar (BMG Labtech) with excitation at 488 nm with 16 nm bandwidth and emission at 527 nm with 10 nm bandwidth and compared to measurements of purified EYFP.

The intracellular concentration of Q40-EYFP in the different strains was determined using the following calculation: amount of EYFP in well/number of animals in the well/cell volume of the tissue. The number of animals in each sample was corrected using the average amount of protein per animal of all samples. Cell volumes were determined on day 1 of adulthood by confocal microscopy using the settings described above. The total fluorescent volumes were quantified using the Voxel Counter plugin in ImageJ. To calculate the average cell volumes, the total fluorescent volumes were then divided by the total number of body wall muscle cells (95) or neurons (302). For this analysis, N = 7 animals were used for body wall muscle cells and N = 9 animals for neurons.

### Kinetic analysis

Data were fitted using the independent nucleation model in Amylofit as described previously (Molenkamp et al., 2021; Sinnige et al., 2021). In brief, the model describes the change of the fraction of cells containing an inclusion, *f(t)*, as:

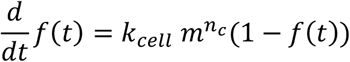

where *k*_*cell*_ is the nucleation rate constant per cell, *m* is the monomer concentration, and *n*_*c*_ is the reaction order of nucleation. Solving the equation and converting the fraction of aggregated cells to the number of aggregated cells, *F(t) = N*_*cells*_ *f(t)*, gives:

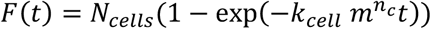

where *N*_*cells*_ is the total number of cells in which nucleation can occur.

To fit inclusion count data for neurons, *N*_*cells*_ was set to 180 for images of the head region and to 302 for images of entire animals. For muscle cells, *N*_*cells*_ was set to 95. *K*_*cell*_ was globally fitted and *n*_*c*_ was fixed to 1.6, unless stated otherwise. *m* was set to 0.766 mM for homozygous neuronal Q40-YFP animals, 0.383 mM for heterozygotes, and 0.751 mM, 0.314 mM, and 0.218 mM for muscle strains A, B, and C, respectively. For HTTex1Q80-YFP, *m* was set to 0.051 mM for homozygotes and 0.0255 mM for heterozygotes. *K*_*cell*_ was converted to the nucleation rate at a fixed monomer concentration by multiplying it with *m*^*nc*^. To compare the nucleation rates in muscle cells and neurons, the nucleation rates were divided by the average cell volume for the respective tissue.

## Data availability

All data are included in the main text and supporting information.

## Author Contributions

Conceptualisation, Writing – original draft, Visualisation: VvS, JT & TS, Investigation, Formal analysis, Writing – Review & editing: All authors, Supervision: JT, MJ & TS.

## Declaration of interest

The authors declare no competing interests.

## Acknowledgement

We thank the Biology Imaging Centre (Utrecht University) for the use of their confocal spinning disk microscope. Some strains were provided by the CGC, which is funded by NIH Office of Research Infrastructure Programs (P40 OD010440). This work was funded by a start-up grant from Utrecht University to TS.

## Notes

### Competing Interest Statement

The authors have declared no competing interest.

### Summary of Updates

Title, introduction and discussion were modified.

## References

Babcock, D. T., & Ganetzky, B. (2015). Transcellular spreading of huntingtin aggregates in the Drosophila brain. Proceedings of the National Academy of Sciences of the United States of America, 112(39), e5427–e5433. 10.1073/pnas.1516217112

Balch, W. E., Morimoto, R. I., Dillin, A., & Kelly, J. W. (2008). Adapting proteostasis for disease intervention. Science, 319(5865), 916–919. 10.1126/science.1141448

Ben-Zvi, A., Miller, E. A., & Morimoto, R. I. (2009). Collapse of proteostasis represents an early molecular event in Caenorhabditis elegans aging. Proceedings of the National Academy of Sciences of the United States of America, 106(35), 14914–14919. 10.1073/pnas.0902882106

Bragg, R. M., Landles, C., Smith, E. J., Osborne, G. F., Mathews, E. W., Cantle, J. P., Bates, G. P., & Carroll, J. B. (2026).Selective targeting of mutant huntingtin intron 1 improves rescue provided by antisense oligonucleotides in Huntington’s disease mice. Science Translational Medicine, 18(841), 702. 10.1126/scitranslmed.adv0702

Brehme, M., Voisine, C., Rolland, T., Wachi, S., Soper, J. H., Zhu, Y., Orton, K., Villella, A., Garza, D., Vidal, M., Ge, H., & Morimoto, R. I. (2014). A chaperome subnetwork safeguards proteostasis in aging and neurodegenerative disease. Cell Reports, 9(3), 1135–1150. 10.1016/j.celrep.2014.09.042

Brenner, S. (1974). The genetics of Caenorhabditis elegans. Genetics, 77(1), 71–94. 10.1093/genetics/77.1.71

Brignull, H. R., Moore, F. E., Tang, S. J., & Morimoto, R. I. (2006). Polyglutamine proteins at the pathogenic threshold display neuron-specific aggregation in a pan-neuronal Caenorhabditis elegans model. Journal of Neuroscience, 26(29), 7597–7606. 10.1523/jneurosci.0990-06.2006

Bunting, E. L., Donaldson, J., Cumming, S. A., Olive, J., Broom, E., Miclăuş, M., Hamilton, J., Tegtmeyer, M., Zhao, H. T., Brenton, J., Lee, W. S., Handsaker, R. E., Li, S., Ford, B., Ryten, M., McCarroll, S. A., Kordasiewicz, H. B., Monckton, D. G., Balmus, G., … Tabrizi, S. J. (2025). Antisense oligonucleotide–mediated MSH3 suppression reduces somatic CAG repeat expansion in Huntington’s disease iPSC–derived striatal neurons. Science Translational Medicine, 17(785), 4600. 10.1126/scitranslmed.adn4600

Chen, S., Ferrone, F. A., & Wetzel, R. (2002). Huntington’s disease age-of-onset linked to polyglutamine aggregation nucleation. Proceedings of the National Academy of Sciences of the United States of America, 99(18), 11884–11889. 10.1073/pnas.182276099

Chiti, F., & Dobson, C. M. (2017). Protein misfolding, amyloid formation, and human disease: A summary of progress over the last decade. Annual Review of Biochemistry, 86, 27–68. 10.1146/annurev-biochem-061516-045115

Cicchetti, F., Lacroix, S., Cisbani, G., Vallières, N., Saint-Pierre, M., St-Amour, I., Tolouei, R., Skepper, J. N., Hauser, R. A., Mantovani, D., Barker, R. A., & Freeman, T. B. (2014). Mutant huntingtin is present in neuronal grafts in huntington disease patients. Annals of Neurology, 76(1), 31–42. 10.1002/ana.24174

Colby, D. W., Cassady, J. P., Lin, G. C., Ingram, V. M., & Wittrup, K. D. (2006). Stochastic kinetics of intracellular huntingtin aggregate formation. Nature Chemical Biology 2006, 2(6), 319–323. 10.1038/nchembio792

Cowan, C. M., & Raymond, L. A. (2006). Selective Neuronal Degeneration in Huntington’s Disease. Current Topics in Developmental Biology, 75, 25–71. 10.1016/s0070-2153(06)75002-5

Crick, S. L., Ruff, K. M., Garai, K., Frieden, C., & Pappu, R. V. (2013). Unmasking the roles of N- and C-terminal flanking sequences from exon 1 of huntingtin as modulators of polyglutamine aggregation. Proceedings of the National Academy of Sciences of the United States of America, 110(50), 20075–20080. 10.1073/pnas.1320626110

DiFiglia, M., Sapp, E., Chase, K. O., Davies, S. W., Bates, G. P., Vonsattel, J. P., & Aronin, N. (1997). Aggregation of huntingtin in neuronal intranuclear inclusions and dystrophic neurites in brain. Science, 277(5334), 1990–1993. 10.1126/science.277.5334.1990

Donaldson, J., Hensman Moss, D., Ciosi, M., Usdin, K., Balmus, G., Monckton, D. G., & Tabrizi, S. J. (2025). Huntington disease: somatic expansion, pathobiology and therapeutics. Nature Reviews Neurology 2025, 22(1), 5–21. 10.1038/s41582-025-01159-7

Donnelly, K. M., Delorenzo, O. R., Zaya, A. D. A., Pisano, G. E., Thu, W. M., Luo, L., Kopito, R. R., & Pearce, M. M. P. (2020). Phagocytic glia are obligatory intermediates in transmission of mutant huntingtin aggregates across neuronal synapses. eLife, 9, e58499. 10.7554/elife.58499

Fawcett, E. M., Hoyt, J. M., Johnson, J. K., & Miller, D. L. (2015). Hypoxia disrupts proteostasis in Caenorhabditis elegans. Aging Cell, 14(1), 92–101. 10.1111/acel.12301

Froehlich, J. J., Rajewsky, N., & Ewald, C. Y. (2021). Estimation of C. elegans cell- and tissue volumes. MicroPublication Biology, 2021. 10.17912/micropub.biology.000345

Fu, H., Hardy, J., & Duff, K. E. (2018). Selective vulnerability in neurodegenerative diseases. Nature Neuroscience, 21(10), 1350. 10.1038/s41593-018-0221-2

Gutekunst, C. A., Li, S. H., Yi, H., Mulroy, J. S., Kuemmerle, S., Jones, R., Rye, D., Ferrante, R. J., Hersch, S. M., & Li, X. J. (1999). Nuclear and Neuropil Aggregates in Huntington’s Disease: Relationship to Neuropathology. Journal of Neuroscience, 19(7), 2522–2534. 10.1523/jneurosci.19-07-02522.1999

Gutierrez-Garcia, R., Koyuncu, S., Hommen, F., Bilican, S., Lee, H. J., Fatima, A., & Vilchez, D. (2023). G3BP1-dependent mechanism suppressing protein aggregation in Huntington’s models and its demise upon stress granule assembly. Human Molecular Genetics, 32(10), 1607–1621. 10.1093/hmg/ddac304

Hartl, F. U., Bracher, A., & Hayer-Hartl, M. (2011). Molecular chaperones in protein folding and proteostasis. Nature 2011, 475(7356), 324–332. 10.1038/nature10317

Herndon, E. S., Hladik, C. L., Shang, P., Burns, D. K., Raisanen, J., & White, C. L. (2009). Neuroanatomic Profile of Polyglutamine Immunoreactivity in Huntington Disease Brains. Journal of Neuropathology & Experimental Neurology, 68(3), 250–261. 10.1097/nen.0b013e318198d320

Herrera, F., Tenreiro, S., Miller-Fleming, L., & Outeiro, T. F. (2011). Visualization of cell-to-cell transmission of mutant huntingtin oligomers. PLoS Currents, 3, RRN1210. 10.1371/currents.rrn1210

Hipp, M. S., Kasturi, P., & Hartl, F. U. (2019). The proteostasis network and its decline in ageing. Nature Reviews Molecular Cell Biology 2019, 20(7), 421–435. 10.1038/s41580-019-0101-y

Hoop, C. L., Lin, H. K., Kar, K., Magyarfalvi, G., Lamley, J. M., Boatz, J. C., Mandal, A., Lewandowski, J. R., Wetzel, R., & Van Der Wel, P. C. A. (2016). Huntingtin exon 1 fibrils feature an interdigitated β-hairpin–based polyglutamine core. Proceedings of the National Academy of Sciences, 113(6), 1546–1551. 10.1073/pnas.1521933113

Kakkar, V., Månsson, C., de Mattos, E. P., Bergink, S., van der Zwaag, M., van Waarde, M. A. W. H., Kloosterhuis, N. J., Melki, R., van Cruchten, R. T. P., Al-Karadaghi, S., Arosio, P., Dobson, C. M., Knowles, T. P. J., Bates, G. P., van Deursen, J.M., Linse, S., van de Sluis, B., Emanuelsson, C., & Kampinga, H. H. (2016). The S/T-Rich Motif in the DNAJB6 Chaperone Delays Polyglutamine Aggregation and the Onset of Disease in a Mouse Model. Molecular Cell, 62(2), 272–283. 10.1016/j.molcel.2016.03.017

Kampmann, M. (2024). Molecular and cellular mechanisms of selective vulnerability in neurodegenerative diseases. Nature Reviews Neuroscience 2024, 25(5), 351–371. 10.1038/s41583-024-00806-0

Kandola, T., Venkatesan, S., Zhang, J., Lerbakken, B. T., Von Schulze, A., Blanck, J. F., Wu, J., Unruh, J. R., Berry, P., Lange, J. J., Box, A. C., Cook, M., Sagui, C., & Halfmann, R. (2023). Pathologic polyglutamine aggregation begins with a self-poisoning polymer crystal. ELife, 12. 10.7554/elife.86939

Kim, D. K., Cho, K. W., Ahn, W. J., Perez-Acuña, D., Jeong, H., Lee, H. J., & Lee, S. J. (2017). Cell-to-cell Transmission of Polyglutamine Aggregates in C. elegans. Experimental Neurobiology, 26(6), 321–328. 10.5607/en.2017.26.6.321

Labbadia, J., & Morimoto, R. I. (2015). The biology of proteostasis in aging and disease. Annual Review of Biochemistry, 84, 435–464. 10.1146/annurev-biochem-060614-033955

Lieberman, A. P., Shakkottai, V. G., & Albin, R. L. (2019). Polyglutamine Repeats in Neurodegenerative Diseases. Annual Review of Pathology: Mechanisms of Disease, 14, 1–27. 10.1146/annurev-pathmechdis-012418-012857

Louros, N., Schymkowitz, J., & Rousseau, F. (2023). Mechanisms and pathology of protein misfolding and aggregation. Nature Reviews Molecular Cell Biology 2023, 24(12), 912–933. 10.1038/s41580-023-00647-2

Månsson, C., Kakkar, V., Monsellier, E., Sourigues, Y., Härmark, J., Kampinga, H. H., Melki, R., & Emanuelsson, C. (2014).DNAJB6 is a peptide-binding chaperone which can suppress amyloid fibrillation of polyglutamine peptides at substoichiometric molar ratios. Cell Stress and Chaperones, 19(2), 227–239. 10.1007/s12192-013-0448-5

Mätlik, K., Baffuto, M., Kus, L., Deshmukh, A. L., Davis, D. A., Paul, M. R., Carroll, T. S., Caron, M. C., Masson, J. Y., Pearson, C. E., & Heintz, N. (2024). Cell-type-specific CAG repeat expansions and toxicity of mutant Huntingtin in human striatum and cerebellum. Nature Genetics 2024, 56(3), 383–394. 10.1038/s41588-024-01653-6

McIntire, S. L., Jorgensen, E., Kaplan, J., & Horvitz, H. R. (1993). The GABAergic nervous system of Caenorhabditis elegans.Nature, 364(6435), 337–341. 10.1038/364337a0

Meisl, G., Hidari, E., Allinson, K., Rittman, T., DeVos, S. L., Sanchez, J. S., Xu, C. K., Duff, K. E., Johnson, K. A., Rowe, J. B., Hyman, B. T., Knowles, T. P. J., & Klenerman, D. (2021). In vivo rate-determining steps of tau seed accumulation in Alzheimer’s disease. Science Advances, 7(44). 10.1126/sciadv.abh1448

Meisl, G., Kirkegaard, J. B., Arosio, P., Michaels, T. C. T., Vendruscolo, M., Dobson, C. M., Linse, S., & Knowles, T. P. J. (2016). Molecular mechanisms of protein aggregation from global fitting of kinetic models. Nature Protocols 2016, 11(2), 252–272. 10.1038/nprot.2016.010

Meisl, G., Knowles, T. P., & Klenerman, D. (2020). The molecular processes underpinning prion-like spreading and seed amplification in protein aggregation. Current Opinion in Neurobiology, 61, 58–64. 10.1016/j.conb.2020.01.010

Meisl, G., Kurt, T., Condado-Morales, I., Bett, C., Sorce, S., Nuvolone, M., Michaels, T. C. T., Heinzer, D., Avar, M., Cohen, S. I. A., Hornemann, S., Aguzzi, A., Dobson, C. M., Sigurdson, C. J., & Knowles, T. P. J. (2021). Scaling analysis reveals the mechanism and rates of prion replication in vivo. Nature Structural & Molecular Biology 2021, 28(4), 365–372. 10.1038/s41594-021-00565-x

Melentijevic, I., Toth, M. L., Arnold, M. L., Guasp, R. J., Harinath, G., Nguyen, K. C., Taub, D., Parker, J. A., Neri, C., Gabel, C.V., Hall, D. H., & Driscoll, M. (2017). C. elegans neurons jettison protein aggregates and mitochondria under neurotoxic stress. Nature 2017, 542(7641), 367–371. 10.1038/nature21362

Michaels, T. C. T., Šarić, A., Habchi, J., Chia, S., Meisl, G., Vendruscolo, M., Dobson, C. M., & Knowles, T. P. J. (2018).Chemical Kinetics for Bridging Molecular Mechanisms and Macroscopic Measurements of Amyloid Fibril Formation. Annual Review of Physical Chemistry, 69, 273–298. 10.1146/annurev-physchem-050317-021322

Molenkamp, J., Den Outer, A., van Schijndel, V., & Sinnige, T. (2021). Monitoring Protein Aggregation Kinetics In Vivo using Automated Inclusion Counting in Caenorhabditis elegans. JoVE (Journal of Visualized Experiments), 2021(178), e63365. 10.3791/63365

Monckton, D. G. (2021). The Contribution of Somatic Expansion of the CAG Repeat to Symptomatic Development in Huntington’s Disease: A Historical Perspective. Journal of Huntington’s Disease, 10(1), 7–33. 10.3233/jhd-200429

Morigaki, R., & Goto, S. (2017). Striatal Vulnerability in Huntington’s Disease: Neuroprotection Versus Neurotoxicity. Brain Sciences, 7(6), 63. 10.3390/brainsci7060063

Morley, J. F., Brignull, H. R., Weyers, J. J., & Morimoto, R. I. (2002). The threshold for polyglutamine-expansion protein aggregation and cellular toxicity is dynamic and influenced by aging in Caenorhabditis elegans. Proceedings of the National Academy of Sciences of the United States of America, 99(16), 10417–10422. 10.1073/pnas.152161099

Nedosekin, D. A., Chen, T. Y., Ayyadevara, S., Zharov, V. P., & Shmookler Reis, R. J. (2021). Label-free photothermal disruption of cytotoxic aggregates rescues pathology in a C. elegans model of Huntington’s disease. Scientific Reports 2021, 11(1), 19732–. 10.1038/s41598-021-98661-x

Neueder, A., Landles, C., Ghosh, R., Howland, D., Myers, R. H., Faull, R. L. M., Tabrizi, S. J., & Bates, G. P. (2017). The pathogenic exon 1 HTT protein is produced by incomplete splicing in Huntington’s disease patients. Scientific Reports 2017, 7(1), 1307–. 10.1038/s41598-017-01510-z

Nollen, E. A. A., Garcia, S. M., Van Haaften, G., Kim, S., Chavez, A., Morimoto, R. I., & Plasterk, R. H. A. (2004). Genome-wide RNA interference screen identifies previously undescribed regulators of polyglutamine aggregation. Proceedings of the National Academy of Sciences of the United States of America, 101(17), 6403–6408. 10.1073/pnas.0307697101

Pearce, M. M. P., Spartz, E. J., Hong, W., Luo, L., & Kopito, R. R. (2015). Prion-like transmission of neuronal huntingtin aggregates to phagocytic glia in the Drosophila brain. Nature Communications 2015, 6(1), 6768–. 10.1038/ncomms7768

Pecho-Vrieseling, E., Rieker, C., Fuchs, S., Bleckmann, D., Esposito, M. S., Botta, P., Goldstein, C., Bernhard, M., Galimberti, I., Müller, M., Lüthi, A., Arber, S., Bouwmeester, T., Van Der Putten, H., & Di Giorgio, F. P. (2014). Transneuronal propagation of mutant huntingtin contributes to non-cell autonomous pathology in neurons. Nature Neuroscience, 17(8), 1064–1072. 10.1038/nn.3761

Peng, C., Trojanowski, J. Q., & Lee, V. M. Y. (2020). Protein transmission in neurodegenerative disease. Nature Reviews Neurology 2020, 16(4), 199–212. 10.1038/s41582-020-0333-7

Poirier, M. A., Li, H., Macosko, J., Cai, S., Amzel, M., & Ross, C. A. (2002). Huntingtin spheroids and protofibrils as precursors in polyglutamine fibrilization. Journal of Biological Chemistry, 277(43), 41032–41037. 10.1074/jbc.m205809200

Preibisch, S., Saalfeld, S., & Tomancak, P. (2009). Globally optimal stitching of tiled 3D microscopic image acquisitions. Bioinformatics, 25(11), 1463–1465. 10.1093/bioinformatics/btp184

Rinauro, D. J., Chiti, F., Vendruscolo, M., & Limbocker, R. (2024). Misfolded protein oligomers: mechanisms of formation, cytotoxic effects, and pharmacological approaches against protein misfolding diseases. Molecular Neurodegeneration, 19(1). 10.1186/s13024-023-00651-2

Sahoo, B., Singer, D., Kodali, R., Zuchner, T., & Wetzel, R. (2014). Aggregation Behavior of Chemically Synthesized, Full-Length Huntingtin Exon1. Biochemistry, 53(24), 3897. 10.1021/bi500300c

Sahtoe, D. D., Andrzejewska, E. A., Han, H. L., Rennella, E., Schneider, M. M., Meisl, G., Ahlrichs, M., Decarreau, J., Nguyen, H., Kang, A., Levine, P., Lamb, M., Li, X., Bera, A. K., Kay, L. E., Knowles, T. P. J., & Baker, D. (2024). Design of amyloidogenic peptide traps. Nature Chemical Biology 2024, 20(8), 981–990. 10.1038/s41589-024-01578-5

Scheres, S. H. W., Ryskeldi-Falcon, B., & Goedert, M. (2023). Molecular pathology of neurodegenerative diseases by cryo-EM of amyloids. Nature 2023, 621(7980), 701–710. 10.1038/s41586-023-06437-2

Silva, M. C., Fox, S., Beam, M., Thakkar, H., Amaral, M. D., & Morimoto, R. I. (2011). A Genetic Screening Strategy Identifies Novel Regulators of the Proteostasis Network. PLOS Genetics, 7(12), e1002438. 10.1371/journal.pgen.1002438

Sinnige, T. (2022). Molecular mechanisms of amyloid formation in living systems. Chemical Science, 13(24), 7080–7097. 10.1039/d2sc01278b

Sinnige, T., Meisl, G., Michaels, T. C. T., Vendruscolo, M., Knowles, T. P. J., & Morimoto, R. I. (2021). Kinetic analysis reveals that independent nucleation events determine the progression of polyglutamine aggregation in C. elegans. Proceedings of the National Academy of Sciences of the United States of America, 118(11). 10.1073/pnas.2021888118

Skeens, A., Siriwardhana, C., Massinople, S. E., Wunder, M. M., Ellis, Z. L., Keith, K. M., Girman, T., Frey, S. L., & Legleiter, J. (2024). The polyglutamine domain is the primary driver of seeding in huntingtin aggregation. PLOS ONE, 19(3), e0298323. 10.1371/journal.pone.0298323

Skibinski, G. A., & Boyd, L. (2012). Ubiquitination is involved in secondary growth, not initial formation of polyglutamine protein aggregates in C. elegans. BMC Cell Biology 2012, 13(1), 10–. 10.1186/1471-2121-13-10

Tabrizi, S. J., Leavitt, B. R., Landwehrmeyer, G. B., Wild, E. J., Saft, C., Barker, R. A., Blair, N. F., Craufurd, D., Priller, J., Rickards, H., Rosser, A., Kordasiewicz, H. B., Czech, C., Swayze, E. E., Norris, D. A., Baumann, T., Gerlach, I., Schobel, S. A., Paz, E., … Lane, R. M. (2019). Targeting Huntingtin Expression in Patients with Huntington’s Disease. New England Journal of Medicine, 380(24), 2307–2316. 10.1056/nejmoa1900907

Thakur, A. K., Jayaraman, M., Mishra, R., Thakur, M., Chellgren, V. M., L Byeon, I. J., Anjum, D. H., Kodali, R., Creamer, T. P., Conway, J. F., M Gronenborn, A., & Wetzel, R. (2009). Polyglutamine disruption of the huntingtin exon 1 N terminus triggers a complex aggregation mechanism. Nature Structural & Molecular Biology, 16(4), 380–389. 10.1038/nsmb.1570

Torricella, F., Tugarinov, V., & Clore, G. M. (2024). Nucleation of Huntingtin Aggregation Proceeds via Conformational Conversion of Pre-Formed, Sparsely-Populated Tetramers. Advanced Science, 11(24), 2309217. 10.1002/advs.202309217

Wagner, A. S., Politi, A. Z., Ast, A., Bravo-Rodriguez, K., Baum, K., Buntru, A., Strempel, N. U., Brusendorf, L., Hänig, C., Boeddrich, A., Plassmann, S., Klockmeier, K., Ramirez-Anguita, J. M., Sanchez-Garcia, E., Wolf, J., & Wanker, E. E. (2018). Self-assembly of Mutant Huntingtin Exon-1 Fragments into Large Complex Fibrillar Structures Involves Nucleated Branching. Journal of Molecular Biology, 430(12), 1725–1744. 10.1016/j.jmb.2018.03.017

Walther, D. M., Kasturi, P., Zheng, M., Pinkert, S., Vecchi, G., Ciryam, P., Morimoto, R. I., Dobson, C. M., Vendruscolo, M., Mann, M., & Hartl, F. U. (2015). Widespread proteome remodeling and aggregation in aging C. elegans. Cell, 161(4), 919–932. 10.1016/j.cell.2015.03.032

Wang, C., Vidal, B., Sural, S., Loer, C., Aguilar, G. R., Merritt, D. M., Toker, I. A., Vogt, M. C., Cros, C. C., & Hobert, O. (2024). A neurotransmitter atlas of C. elegans males and hermaphrodites. ELife, 13. 10.7554/elife.95402

Zhang, Y., Han, H. L., Ortigosa-Pascual, L., Miles, U. Z., Wei, J., Berkowicz, L., Snow, F., Tu, D., Meisl, G., Sansaria, S., Nott, T. J., Laman, H., Halfmann, R., McShan, A. C., Sahtoe, D. D., & Knowles, T. P. J. (2026). State-selective inhibition of alpha-synuclein aggregation by de novo oligomer-binding proteins. BioRxiv, 2026.07.13.738012. 10.64898/2026.07.13.738012

