## Supplementary information for "Polyglutamine aggregation in the *C. elegans* nervous system is driven by stochastic nucleation events"

**Table S1.** Cell volumes of *C. elegans* neurons versus body wall muscle cells

| Dataset | Total volume (pL) | Volume per cell (pL) | Reference |
|---|---|---|---|
| Neurons | 39.4 | 0.13 | This study |
| Neurons (Blender) | 12.7 | 0.04 | (Froehlich et al,. 2021) |
| Neurons (Mesh Volume Calculator) | 19.9 | 0.07 | (Froehlich et al,. 2021) |
| Muscle | 172 | 1.81 | This study |
| Muscle | 99 | 1.04 | (Sinnige et al,. 2021) |
| Muscle (Blender) | 166 | 1.75 | (Froehlich et al,. 2021) |
| Muscle (Mesh Volume Calculator) | 182 | 1.91 | (Froehlich et al,. 2021) |

**Table S2.** Nucleation rates of HTTex1Q80-YFP compared to Q40-YFP in neurons

| **Dataset** | **$K_{cell}$ ($M^{-1.6}$ $d^{-1}$)** | **Nucleation rate at 1 mM ($d^{-1}$ $cell^{-1}$)** | **Nucleation rate at 1 mM ($d^{-1}$ $pL^{-1}$)** |
|---|---|---|---|
| Q40-YFP | $1.04 \times 10^3$ | 0.016 | 0.13 |
| HTTex1Q80-YFP | $4.66 \times 10^4$ | 0.739 | 5.68 |

**Table S3.** *C. elegans* strains used in this study

| Strain name | Genotype | Source |
|---|---|---|
| AM52 | rmIs182 [p*F25B3.3*::Q0::YFP] | Morimoto lab |
| AM101 | rmIs110 [p*F25B3.3*::Q40::YFP] | CGC |
| AM138 | rmIs130 [p*unc-54*::Q24::YFP] | CGC |
| AM716 | rmIs284 [p*F25B3.3*::Q67::YFP] | Morimoto lab |
| AM1228 | rmIs404 [p*unc-54*::Q40::YFP] line A | Morimoto lab |
| AM1230 | rmIs404 [p*unc-54*::Q40::YFP] line B | Morimoto lab |
| AM1231 | rmIs404 [p*unc-54*::Q40::YFP] line C | Morimoto lab |
| TSW49 | mbbIs18[p*rab-3*::FLAG::mKate2::DNAJB6::*let-858* 3'UTR] | This study |
| TSW69 | mbbIs12[p*unc-54*::FLAG::mKate2::DNAJB6::*let-858* 3'UTR] | This study |
| TSW82 | rmIs404 [p*unc-54*::Q40::YFP] line A; mbbIs12[p*unc-54*::FLAG::mKate2::DNAJB6::*let-858* 3'UTR] (cross of TSW49 x AM101) | This study |
| TSW83 | rmIs404 [p*unc-54*::Q40::YFP] line B; mbbIs12[p*unc-54*::FLAG::mKate2::DNAJB6::*let-858* 3'UTR] (cross of TSW69 x AM1230) | This study |
| TSW84 | rmIs404 [p*unc-54*::Q40::YFP] line C; mbbIs12[p*unc-54*::FLAG::mKate2::DNAJB6::*let-858* 3'UTR] (cross of TSW69 x AM1231) | This study |
| TSW85 | rmIs110 [p*F25B3.3*::Q40::YFP]; mbbIs18[p*rab-3*::FLAG::mKate::DNAJB6::*let-858* 3'UTR] (cross of TSW49 x AM101) | This study |
| TSW86 | rmIs284[p*F25B3.3*::Q67::YFP] ; mbbIs18[p*rab-3*::FLAG::mKate::DNAJB6::*let-858* 3'UTR] (cross of TSW49 x AM716) | This study |
| TSW65 | mbbIs15[p*rgef-1*::Htt exon 1 Q20::mEYFP::*let-858* 3'UTR] | This study |
| TSW66 | mbbIs16[p*rgef-1*::Htt exon 1 Q80::mEYFP::*let-858* 3'UTR] | This study |
| TSW93 | mbbIs16[p*rgef-1*::Htt exon 1 Q80::mEYFP::*let-858* 3'UTR]; mbbEx10[p*rab-3*::FLAG::mKate::DNAJB6::*let-858* 3'UTR] | This study |

**Table S4.** Oligonucleotides used in this study

| Primer | Primer sequence |
|---|---|
| p*unc-54* forward primer | AACATATCCAGTCACTATGGCTGCAGTGAGTATTTTCGG |
| p*unc-54* reverse primer | CATTTTTCTGAAAAGCCTGCTACGTCTCGCTTCTTTCAAATGGTT |
| p*rgef-1* forward primer | AACATATCCAGTCACTATGGATCACAAATAAGGAATAAGCGACG |
| p*rgef-1* reverse primer | CATTTTTCTGAAAAGCCTGCTACGTACGAACGATTGAGCAGAAGG |
| p*rab-3* forward primer | AACATATCCAGTCACTATGGCATATTTTTTGACGACGACGAC |
| p*rab-3* reverse primer | CATTTTTCTGAAAAGCCTGCTACGTCTGAAAATAGGGCTACTGTAGATTTA |
| *let-858* 3'UTR forward primer | CGATCGTGAAGTGGAATCGGATGA |
| *let-858* 3'UTR reverse primer | GAGAAAATACCGCATCAGGCATACGGATTCGCATTTGCCAAG |
| mKate2 forward primer | AGGCTTTTCAGAAAAATGGACTACAAAGACGATGACGACAAGATGGTCTCCGAGCTCATT |
| mKate2 reverse primer | GCCCACTTTGTACAAGAAAGCTGGGTCACGGTGTCCGAGCTTGGATG |
| mEYFP forward primer | GACCCAGCTTTCTTGTACAAAGTGGGCATGGTGAGCAAGGGCGAGGA |
| mEYFP reverse primer | GATTCCACTTCACGATCGCTTGTCGTCATCGTCTTTGTAGTCTTACTTGTACAGCTCGTC |
| HTTex1 forward primer | ACGTAGCAGGCTTTTCAGAAAAATGGCAACGCTGGAAAAACTG |
| HTTex1 reverse primer | GCCCACTTTGTACAAGAAAGCTGGGTCCGGGCGATGCAGCGGTTC |

**Table S5.** Plasmids used in this study

| Plasmid name | Description |
|---|---|
| pJT07 | EYFP bacterial expression vector |
| pJT18 | p*rab-3*::1xFLAG::mKate2:hDNAJB6::*let-858* 3'UTR |
| pKB06 | p*unc-54*::1xFLAG::mKate2:hDNAJB6::*let-858* 3'UTR |
| pDS01 | p*rgef-1*::HTTexon1-Q20::mEYFP::*let-858* 3'UTR |
| pDS03 | p*rgef-1*::HTTexon1-Q80::mEYFP::*let-858* 3'UTR |

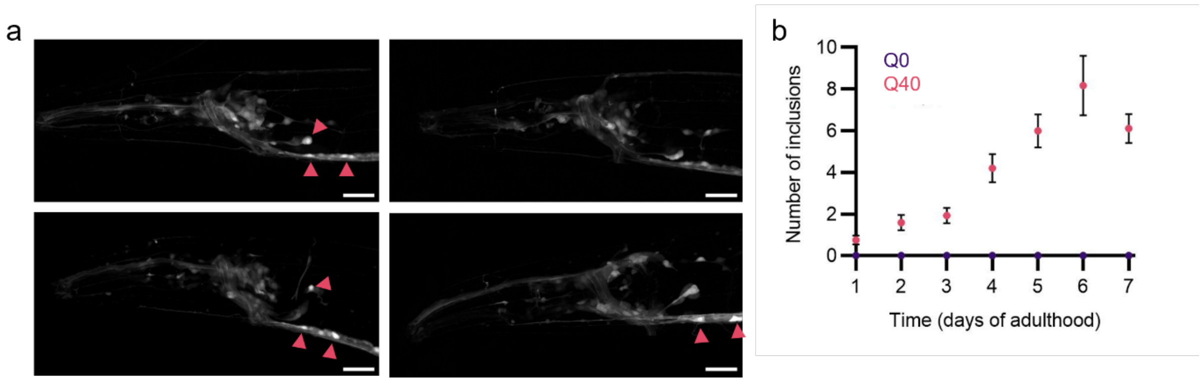


**Figure S1.** Inclusions are absent in control animals. **a** Confocal images of *C. elegans* head region of Q0-YFP animals imaged at day 5 of adulthood. Arrowheads indicate bright cell bodies. **b** Quantification of inclusion numbers in the head region over time in animals expressing Q40-YFP versus Q0-YFP, using the same imaging and analysis settings. Data represent mean ± SEM (N = 20 animals per condition).

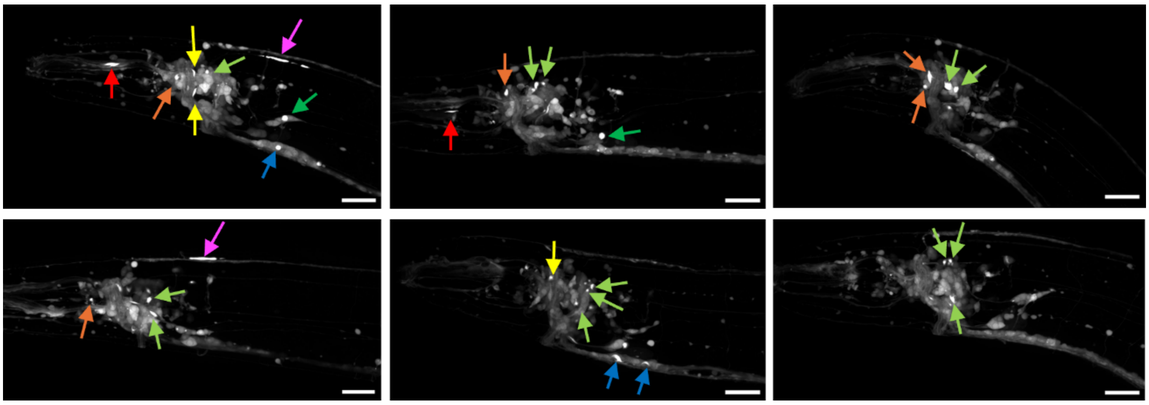

**Figure S2.** Q40-YFP inclusions localise to different parts of the nervous system. Shown are confocal images of *C. elegans* head region of Q40-YFP animals. Arrows indicate inclusions in the dendrites of sensory neurons (red), anterior ganglion (orange), nerve ring (yellow), lateral ganglion (green), posterior head neurons (dark green) ventral ganglion (blue), dorsal cord (magenta). Scale bars: 20 μm.

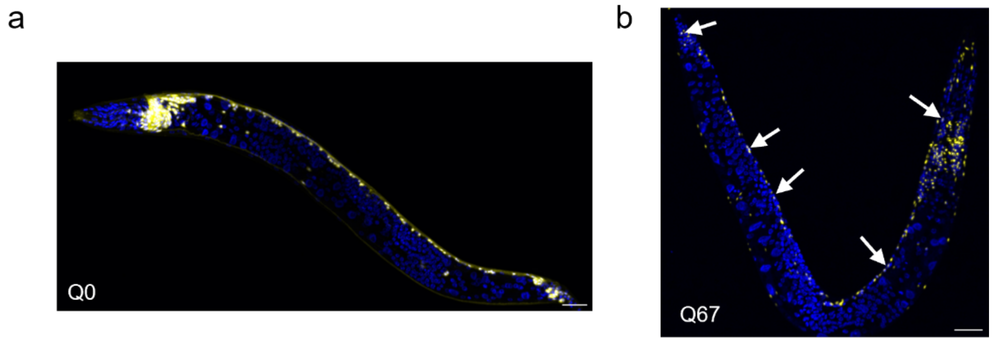


**Figure S3**. Q67-YFP localizes to both the nucleus and cytoplasm in all neurons of the animal. **a** Confocal image of a fixed L4-staged animal expressing Q0-YFP (yellow) stained with DAPI (blue). **b** Confocal image of a fixed L4-staged animal expressing Q67-YFP (yellow) stained with DAPI (blue). Arrows indicate overlap between Q67-YFP and DAPI (visible as white spots). Scale bars: 20 μm.

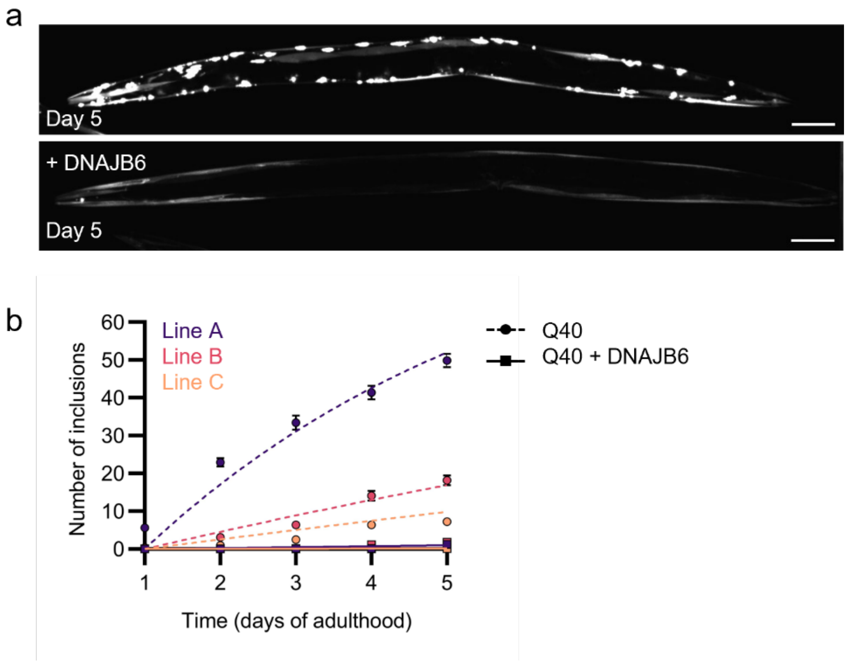


**Figure S4**. DNAJB6 inhibits Q40-YFP aggregation in body wall muscle cells. **a** Confocal images of day 5 animals expressing Q40-YFP (line B) in body wall muscle cells under control conditions and upon overexpression of human DNAJB6. Scale bars: 50 µm. **b** Kinetics of Q40-YFP aggregation upon DNAJB6 overexpression compared to controls. Inclusions were quantified over time in animals expressing different concentrations of Q40-YFP (lines A-C). Solid and dashed lines represent fits to the independent nucleation model for DNAJB6 overexpression and control animals respectively. Data represent mean ± SEM (N = 20 animals per condition).

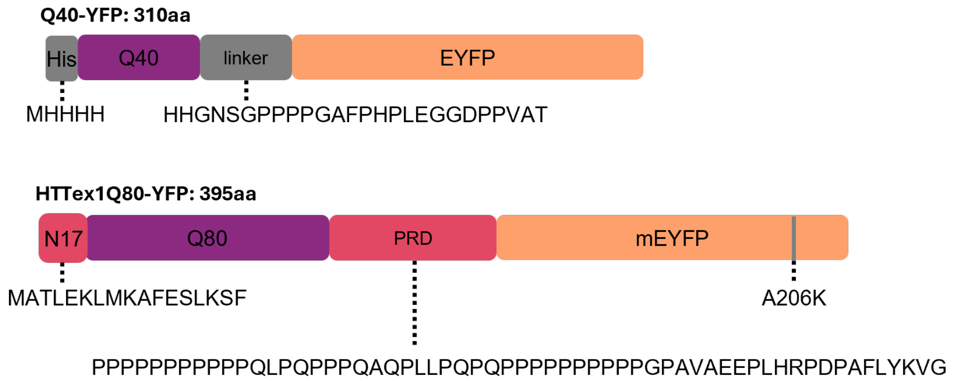


**Figure S5.** Schematic illustration of the Q40-YFP and HTTex1Q80-YFP constructs. Q40-YFP contains an N-terminal His-tag, a 40-residue polyglutamine (Q40) tract, and a short linker followed by EYFP. HTTex1Q80-YFP contains the N17 domain of huntingtin, a 80-residue polyglutamine (Q80) tract, the proline-rich domain (PRD), and monomeric EYFP (mEYFP), which carries the A206K monomerizing mutation. Amino acid sequences of the linkers and flanking regions are shown below each schematic.

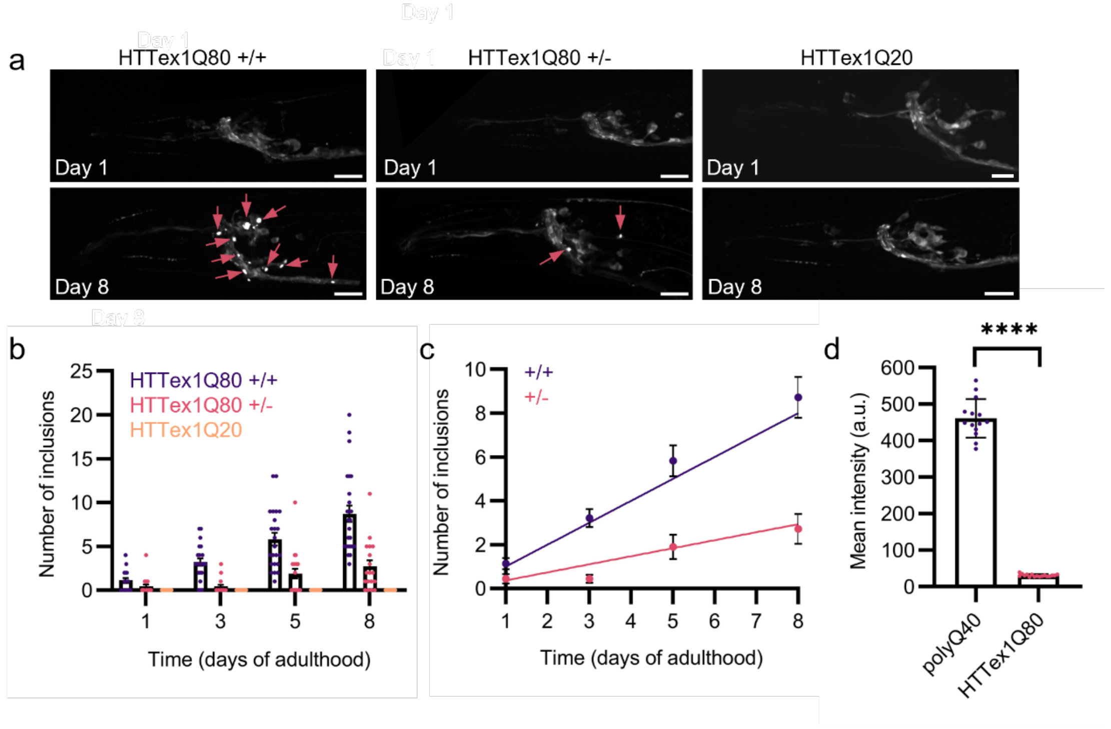


**Figure S6.** Characterisation of *C. elegans* expressing HTTex1. **a** Confocal images of homozygous animals expressing HTTex1Q80-YFP (left), heterozygous animals expressing HTTex1Q80-YFP (middle), and animals expressing HTTex1Q20-YFP (right) on day 1 and 8 of adulthood. Scale bars: 20 µm. **b** Quantification of inclusion numbers in the head region of animals expressing HTTex1Q80-YFP versus HTTex1Q20-YFP, showing that no inclusions are detected in the latter strain. Data represent mean ± SEM (N = 18 - 25 animals per condition). **c** Inclusion numbers for animals homozygous versus heterozygous for HTTex1Q80-YFP, fitted to a model of independent nucleation events occurring in individual cells. The maximum number of cells was set to 180 and $n_c$ was set to 1.6. $K_{cell}$ was fitted globally. Data represent mean ± SEM (N = 18 - 25 animals per condition). **d** Quantification of the mean integrated density in animals expressing Q40-YFP and HTTex1Q80-YFP on day 1 of adulthood. Data represent mean ± SEM (N = 14-17 animals per condition). $p < 0.0001$ (****).

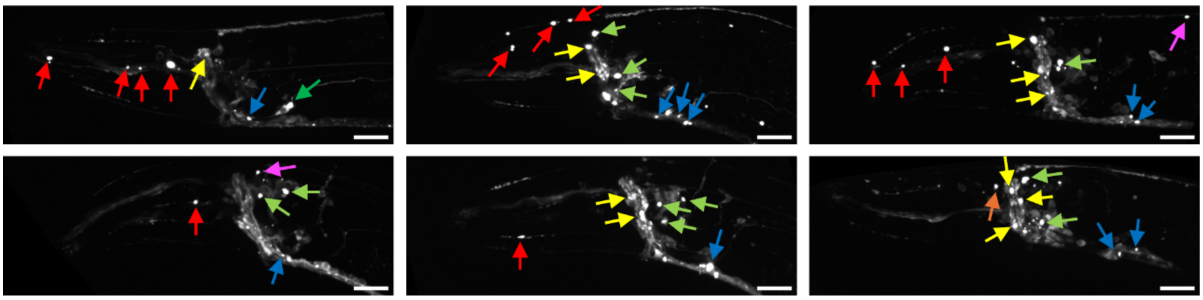

**Figure S7.** HTTEx1Q80-YFP inclusions localise to different parts of the nervous system. Shown are confocal images of *C. elegans* head region of HTTex1Q80-YFP animals on day 8 of adulthood. Arrows indicate inclusions in the dendrites of sensory neurons (red), anterior ganglion (orange), nerve ring (yellow), lateral ganglion (green), posterior head neurons (dark green) ventral ganglion (blue), dorsal cord (magenta). Scale bars: 20 µm.

### References

Froehlich, J. J., Rajewsky, N., & Ewald, C. Y. (2021). Estimation of *C. elegans* cell- and tissue volumes. *MicroPublication Biology*, *2021*. https://doi.org/10.17912/micropub.biology.000345

Sinnige, T., Meisl, G., Michaels, T. C. T., Vendruscolo, M., Knowles, T. P. J., & Morimoto, R. I. (2021). Kinetic analysis reveals that independent nucleation events determine the progression of polyglutamine aggregation in *C. elegans*. *Proceedings of the National Academy of Sciences of the United States of America*, *118*(11). https://doi.org/10.1073/pnas.2021888118
